# Long-term, multi-event surprise enhances autobiographical memory

**DOI:** 10.1101/2022.11.27.517985

**Authors:** James W. Antony, Jacob Van Dam, Jarett R. Massey, Alexander J. Barnett, Kelly A. Bennion

## Abstract

Neurobiological and psychological models of learning emphasize the importance of prediction errors (surprises) for memory formation. These effects have focused on memory for information surrounding a momentary surprising event; however, it is less clear whether surprise that unfolds across multiple events and timescales impacts memory. We asked basketball fans about their most positive and negative autobiographical memories of individual plays, games, and seasons, allowing surprise measurements spanning seconds, hours, and months. We used advanced analytics on National Basketball Association play-by-play data and betting odds spanning 17 seasons, >22K games, >5.6M plays to compute and align the estimated surprise values of each memory. We found that surprising events biased positive memories on the scale of seconds and months and negative memories across all three timescales. Critically, game and season memories could not be explained by surprise at shorter timescales, suggesting that long-term, multi-event surprise influences memory. These results expand the role of surprise in models of learning and reinforce its relevance in real-world domains.

## Introduction

Predictions about upcoming events – and their eventual outcomes – have a substantial effect on learning and memory^1–7^. A wealth of research has demonstrated that momentary prediction errors, or surprises, are particularly well-remembered^1,2,7–12^ and that an array of physiological and mental processes seem to underpin these memory benefits. For instance, surprise engages the dopaminergic and serotonergic midbrain systems^7,10,13–17,^ alters and enhances hippocampal activity^16,18–24^, increases pupil dilation^7,25–29^, and enhances perceptual^13,16,30,31^ and attentional processing^8,32–39^ to the surprising stimulus. Surprises can be further differentiated into events that are better than expected for the agent because they signal reward (+ reward prediction error, or + signed surprise) and others that are worse than expected (-signed surprise)^40–45^. The physiological response to these events is critical for learning and updating values of stimuli in the service of the optimal behavior to approach rewards and avoid punishments^41^. With regard to memory, outcomes with + signed surprise are better remembered in some paradigms^3,40,46,47^, whereas in others, the magnitude, rather than the direction of surprise (or unsigned surprise) drives memory^1,2,7,11,40^.

The prevailing focus in these studies is how surprise affects learning and memory for momentary events on the scale of seconds; however, it is clear that humans can predict outcomes well beyond the upcoming moment^48^. A very non-exhaustive list of things we can make predictions about beyond the next event include a series of upcoming stimuli^49,50^, the linguistic content of not just upcoming words^51–53^ but paragraphs^54,55^, attributes about oneself months and years into the future^56–59^, political elections months and years into the future^60^, and the winners of upcoming sports games^61,62^ and championships^63^. Given that we commonly make long-term predictions and that surprise drives memory, it is worth asking whether longterm predictions resulting in errors (long-term surprises) also drive memory.

The problem of long-term predictions is intriguing in domains like politics or sports that feature probabilistic updates about a single outcome over long stretches of time. As humans can represent information at multiple timescales of granularity, it is unclear which timescale should be operative at any given time, whether errors accumulate over time, or whether each of these variables impact memory. Consider the following example: Imagine you predicted in late January 2020 that Bernie Sanders would become the Democratic nominee for US president, which was ∼ 25% likely per 538. In this hypothetical scenario, imagine that throughout February and March 2020, Sanders won state after state, and therefore Sanders’ likelihood inched upwards every few days until it reached 98% by April 1. Had Sanders won one more state on April 7 and secured the nomination (100%), would it be surprising? The answer depends on the timescale. If judging from April 1, no: with only a 2% surprise, the outcome was essentially a foregone conclusion. However, if judging from late January, at 75% surprise, the answer is undoubtedly yes. Importantly, these kinds of events are both long-term (the interval between an initial prediction and final outcome spans more than a single moment) and composed of multiple sub-events (there are numerous updates to predictions before that outcome).

Here our primary focus was to determine whether surprise occurring across multiple timescales and events affects memory. To do so, we framed our questions around events spanning multiple moments – where the events could still be surprising as a collective even if they were not individually. For this, we asked basketball fans about their most positive and negative memories of entire basketball games and seasons. To link these results with the literature on momentary surprise, we also asked fans about individual plays, allowing us to replicate findings in this literature and extend them to real-world memories (see also^64,65^). We chose the sports domain for three main reasons. First, recent explosions in publicly available sports data (e.g.,^66^) allowed us to precisely quantify predictions about the likelihoods that a given team would win a game or championship. Using such predictions, we were able to form estimates of surprise following the outcome of a play, game, or season. Second, sports contain a natural hierarchy of timescales. In the National Basketball Association (NBA), which is the focus here, the broadest scale over which teams compete is the season, and seasons consist of collections of games, which consist of collections of plays. Therefore, we could find and situate the context of subject memories within a multi-scale predictive framework with respect to game and championship likelihoods. Third, affiliating with sports teams often provides fans a strong sense of identity^67,^ and as a result, sports memories can be highly arousing and vivid^68,69^. Emotional events are recollected more vividly^70–73^, more often^74–76^, and for longer intervals than neutral events^74,77,78^, and they also often serve as the basis for experiments surrounding highly vivid, “flashbulb” memories of public events^79–82^. We therefore anticipated that subjects would be able to readily access these memories. We predicted that surprise would play an outsize role in memory across all three timescales and, critically, that the contributions of surprise at the longer timescales could not be attributed to surprise at shorter scales.

## Results

### Characterizing subject responses

Subjects (N=122, 34 female) took a survey asking them about their most positive and negative memories of individual plays, games, and seasons as basketball fans (Figure 1A). We asked them to include as much detail as possible, especially emphasizing that we would like to specifically find each memory via later internet search. Our goal was to precisely identify the play, game, or season from which subject memories occurred so we could quantify their surprise-related attributes. We intentionally left the question open-ended and avoided implying that they should use surprise as a heuristic. It was slightly more difficult to identify precise plays and games than seasons: in total, we identified 70 positive NBA plays, 74 negative plays, 77 positive games, 74 negative games, 96 positive seasons, and 83 negative seasons.

**Figure 1:**
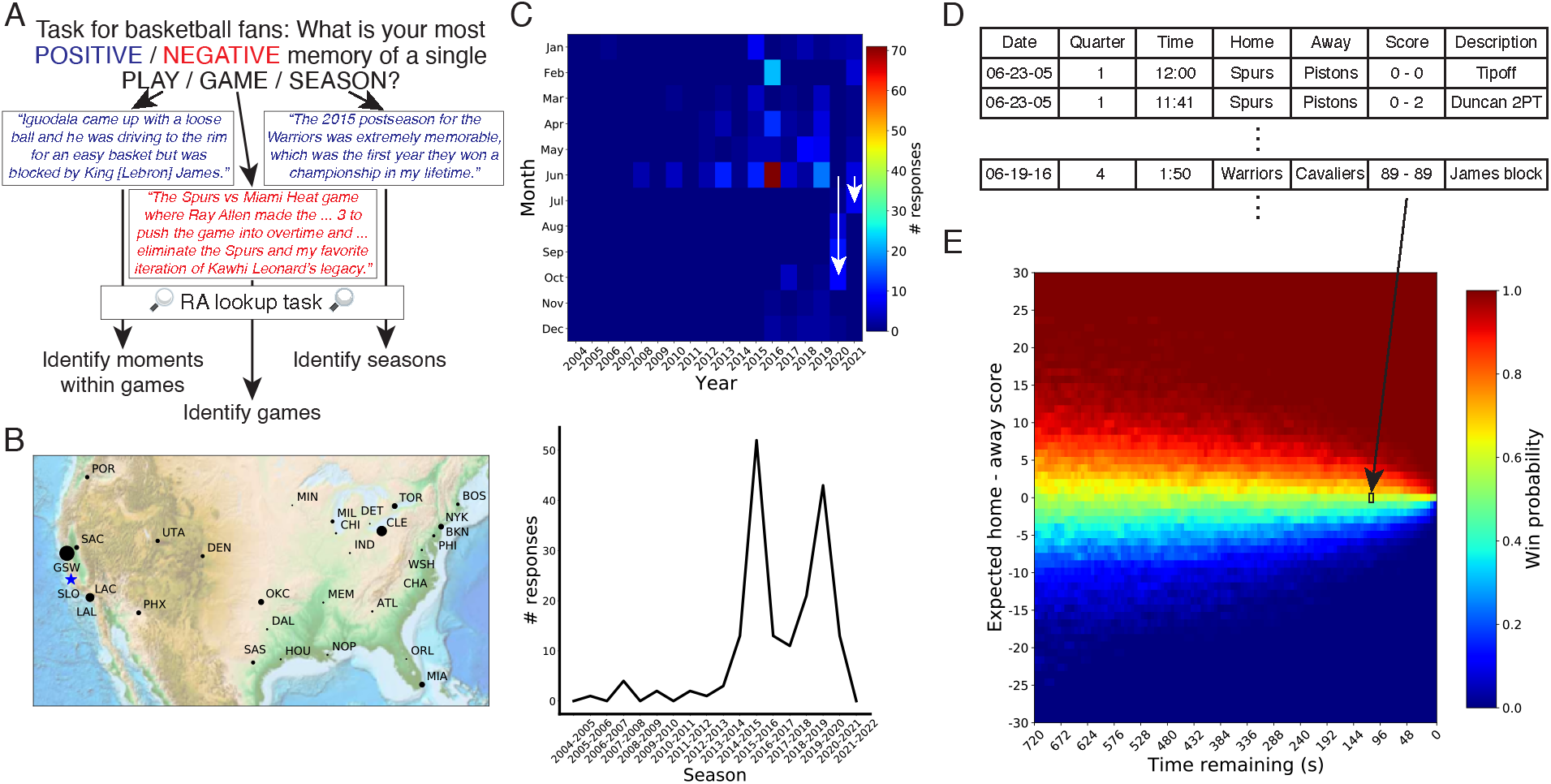
Memory task, basic response characteristics, and analytical approach. (A) Subjects were asked to freely respond to prompts about their most positive and negative memories of plays, games, and seasons as basketball fans. We then linked these responses to specific teams and times across multiple time scales (moments in games, games within seasons, seasons). (B-C) We characterized the response frequencies by the primary NBA team’s geographical location for plays, games, and seasons (B) as well as their temporal distributions by month and year of play and game responses (C, top) and starting year of season responses (C, bottom). Due to the Covid-19 pandemic, the end of the NBA season was shifted four months in 2020 and one month in 2021 (white arrows in C). (D-E) We imported play-by-play data for all games across 17 seasons (2004-05 to 2020-2021) (D) to find game context of all subject responses and to train a win probability model (E). The win probability model computed the likelihood of winning the game given each game context, including the strength of the two teams and the current score (which were combined into an adjusted score), the time remaining in the game, and the team in possession of the ball (only ‘Home’ shown in E). For instance, the play from (A) in which Lebron James blocked a shot corresponds to the win probability in the black square in (F).

Twenty-nine of the thirty NBA teams served as the primary team of interest in subject responses (Figure 1B; Figure S1). We conducted the study in a location that was >320 km (200 miles) from the nearest NBA team stadium in order to reduce the bias towards a single fanbase, though there remained a bias towards the nearest teams (responses across all types, the San Francisco-area-based Golden State Warriors: 261; Los Angeles Lakers: 83) (Figure 1B). Memories for plays and games were biased towards recent years (after 2013) and towards the end of the postseason, which typically occurred in June but was moved to October in 2020 and July in 2021 for Covid-19 pandemic rescheduling (Figure 1C, top). Memories for seasons were similarly biased towards recent years (Fig 1C, bottom). As data were collected from February until November 2021, we also plotted the age of the memories between the study and the date of their reported memories, which showed a similar bias (Figure S2).

### Momentary surprise drove memory for positive and negative plays

We began our investigations by examining surprise for positive and negative plays, which resembles the momentary surprise studied in many human memory paradigms (e.g.,^11^) and in traditional reinforcement learning paradigms (e.g.,^43^). Our main analytical approach for play and game memories relied on scraping the NBA API for play-by-play data (Figure 1D). These data included >5.6M plays from >22K games between 2004-2021. Our derivation of momentary surprise was based on the primary outcome variable of relevance to sports fans: which team wins a given game. One can conceptualize watching a basketball game as navigating a state space of predictions about the eventual winner that becomes updated with each change in game context (Figure 1E; Figure S3). Four factors influenced win probability in our model: the score difference between the two teams (oriented as positive for the home team), the relative strength of the teams, the amount of game time remaining, and the team with possession of the ball. Context changes included scores, timeouts, and three types of plays that potentially change possession (turnovers, rebounds, and tip-offs). We then computed surprise as the derivative in the win probability time course across these changes in game context (Figure 2A). This construct was later split into an unsigned value using the absolute value of this derivative and a signed value that was oriented positively for each subject’s preferred team and negatively for their non-preferred team. Importantly, we validated that the surprise metrics from all positive and negative plays based on our algorithm corresponded tightly with those from an [expert website](http://www.inpredictable.com) (Figure S4B).

**Figure 2:**
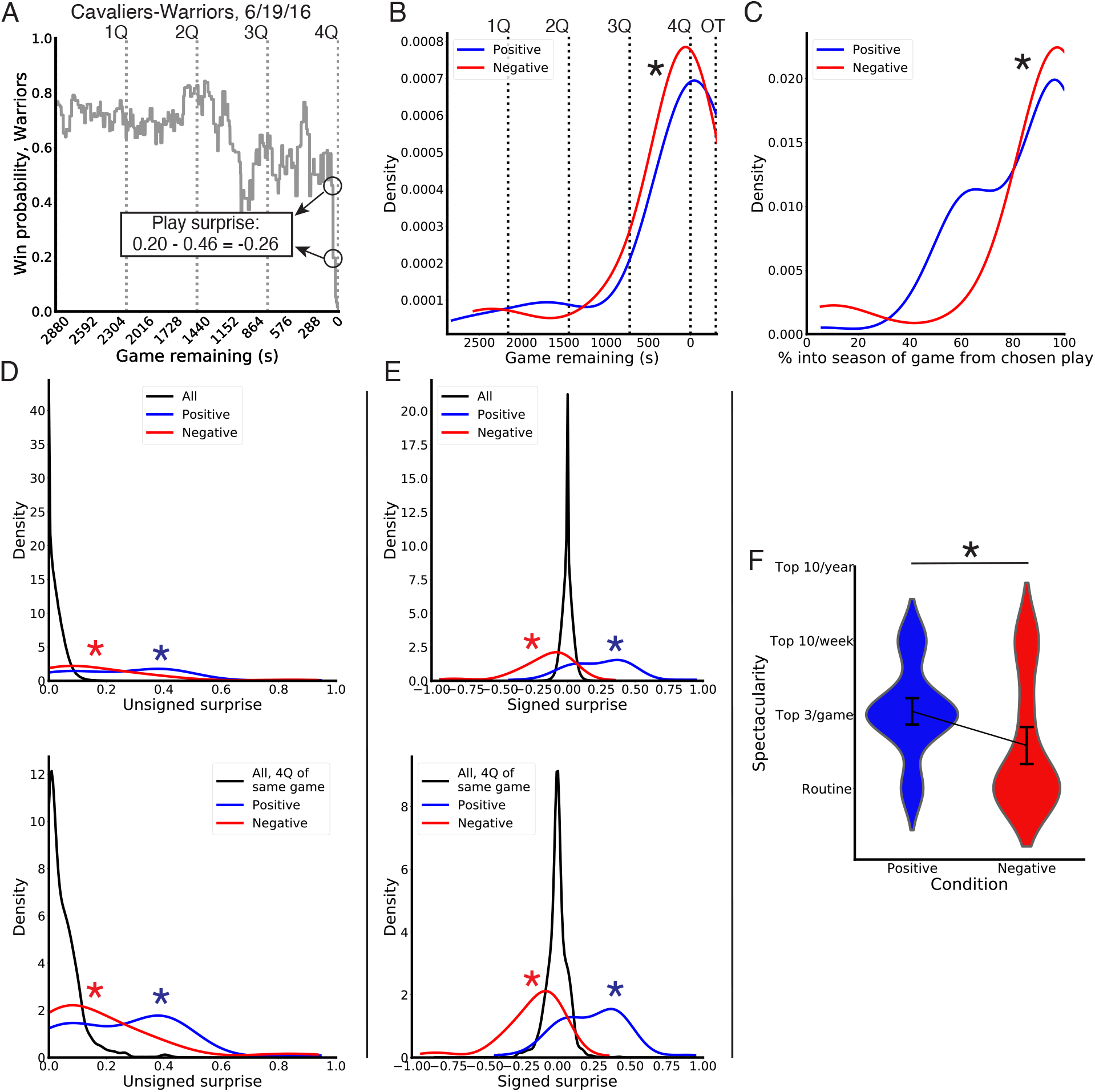
Surprise predicts memory for positive and negative plays. (A) We computed play surprise as the difference in win probabilities across successive moments, shown here using Game 7 from the 2016 NBA Finals between the Cleveland Cavaliers and Golden State Warriors. We computed an unsigned version of this metric using the absolute value and a signed version of this metric by considering this value with respect to the subject’s preferred team (e.g., positive for a preferred outcome). (B) Positive and negative responses were strongly biased towards the end of games. 1-4Q: 1^st^-4^th^ quarter; OT: 1^st^ overtime. (C) Positive and negative responses were also strongly biased towards the end of the corresponding team’s season. (D) Positive and negative responses corresponded with moments of higher unsigned surprise than a null distribution of all basketball plays (top) and a more focused null distribution of the 4^th^ quarter of the games chosen by subjects (bottom). (E) Positive and negative responses corresponded to higher and lower signed surprise, respectively, than null distributions of all basketball plays (top) and the 4^th^ quarter of the games chosen by subjects (bottom). (F) The spectacularity of a given play, which assesses an alternative way in which a basketball play can be surprising that is unrelated to win probability, was higher for positive than negative play responses. *: *p* < 0.01

Before characterizing surprise, we investigated when these plays occurred within games and within seasons. Both positive and negative plays tended to come from the very end of the 4^th^ quarter of games and into overtime periods, and positive plays tended to have less time remaining in the game than negative plays [positive: median = 5 s, *N* = 70; negative: median = 53 s, *N* = 74; Mann-Whitney *U* = 1904.5, *p* = 0.006] (Figure 2B). Interestingly, examining the mean and standard deviation of surprise across all games revealed that mean surprise went down modestly throughout the average game, accounting for the large number of games in which the outcome is all but certain by the end (“blowouts”) (Figure S5A). However, standard deviation of surprise increased exponentially towards the end of the game, suggesting the final moments offer the largest potential for surprise (Figure S5B). We also calculated the percentile of the game from which the play occurred relative to the chosen team’s season. For example, the first game of the season was in the 0th percentile and the last game was in the 100th percentile. Similar to how responses tended to come from the final months (Figure 1C), these responses tended to come from the very end of the primary team’s season. In this case, negative plays tended to come closer to the end of a team’s season than positive plays [median percentile, positive: 89.3th; negative: 99.0th; Mann-Whitney *U* = 1922, *p* = 0.006] (Figure 2C).

Based on prior results showing that momentary surprise influences memory^1,2,7,11,40^, we predicted that memories for positive and negative plays would preferentially be selected from a subset of the most surprising plays. We first tested this by examining whether unsigned surprises from the chosen play memories were higher than a null distribution of all plays from all games. It should be noted that the null distribution was non-normal – most plays had little relevance on the eventual outcome: 32.4% and 78.6% of plays had *<*0.01 and *<*0.05 surprise, respectively, whereas only 0.1% had >0.25 surprise. Subject play memories were significantly greater than this null distribution overall [median, positive: 0.31; negative: 0.11, null: 0.019; positive vs. null Mann-Whitney *U* = 3.4*10^8^, *p* < 0.001; negative vs. null Mann-Whitney *U* = 3.2*10^8^, *p* < 0.001; positive vs. negative Mann-Whitney *U* = 3372, *p* = 0.002], with 78.5% and 51.4% of positive plays and 71.6% and 17.6% of negative plays having >0.05 and >0.25 surprise (Figure 2D, top). We next reasoned that sports fans have limited time and likely do not watch every game or part of every game. Instead, in order to maximize their experience of surprise^83^, which acts as a reward in itself in low-stakes contexts like sports games^7,62,84–88^, they may prioritize their fandom by watching select games or parts of games. If this were the case, a null distribution of plays that is more representative of the plays actually experienced by subjects would be those from the games the subjects reported, or even just plays from the 4^th^ (final) quarter of those games. Positive and negative play surprise was higher than the null distribution from only the games that subjects reported (null median: 0.025; Mann-Whitney *U >* 1*10^6^, *p* < 0.001, both) and also higher than plays from the 4^th^ quarters (null median: 0.036; Mann-Whitney *U* > 2.5*10^5^, *p* < 0.001, both) (Figure 2D, bottom).

We next asked whether these memories represented particularly positive or negative signed surprises with respect to the subject’s preferred team^7,43,89^. For our null distribution, we used a double distribution of the same plays, as if assuming a hypothetical fan might cheer for each team in each game. We again used all plays from all games, all plays from the chosen games, and all plays from the 4^th^ quarter of the chosen games. In all cases, positive memories were more positive and negative memories were more negative than the null distributions (median, positive: 0.29; negative: -0.11, null in all cases: 0.0; Mann-Whitney *U* > 6.5*10^5^, *p* < 0.001) (Figure 2E). Therefore, subjects tended to report plays with exceptionally good or bad win probability swings for their preferred teams. In sum, the influence of unsigned and signed surprises on memory for plays replicates prior findings on the influence of momentary surprise on memory^7,40^ and extends them into real-world domains^64,65^.

Finally, we also considered that our operationalization of surprise as win probability change is only one dimension along which basketball can be surprising^7^. Rather, one might consider a surprising play in terms of the way it unfolds against a lifetime of viewing other plays, such as a spectacular or improbable shot. Since viewing feats of athleticism may be one of the primary reasons one watches sports, we hypothesized that such plays would be more likely to be categorized as positive, though we acknowledge the alternative possibility that an improbable shot from an opposing team may also “sting” more deeply. We operationalized this type of surprise as the ‘spectacularity’ of a given play, which we separately rated without regard for the context of the game. We rated these plays as (1) a routine play like a layup or baseline jump shot, (2) a tough play like a long shot or a fadeaway jumpshot that might be considered in the top 3 in a given game, (3) a particularly athletic play that might be featured on a “Top 10 plays of the day” news segment across all sports, such as a half-court shot or behind-the-basket, changing-hands-in-mid-air layup, or (4) a truly sensational play that might be considered on a year-end “Top 10 plays of the year” segment like a full-court shot or a famous dunk by Michael Jordan off of a missed free throw attempt. We found that positive plays had higher spectacularity than negative plays [positive: median = 2, mean = 2.0, *N* = 45; negative: median = 1, mean = 1.6, *N* = 48; Kolmogorov-Smirnov (KS) statistic = 0.49, *p* < 0.001], indicating that spectacularity was another dimension influencing subject memories and that it was higher for positive than negative memories (Figure 2F).

### Long-term surprise influenced negative memories for games

Next, we aimed to determine whether surprise on a longer timescale – lasting beyond a single moment – would also influence memory. For this analysis, we considered two long-term surprise metrics calculated over the course of games. The first was full-game surprise, or the difference between the pre-game win probability (i.e., with 2880 seconds remaining) and the final result. The second metric was within-game surprise, or the difference between the lowest probability for the eventual winner and the final result (Figure 3A). This metric captured the familiar concept of a “comeback” that looks at how maximally wrong one’s predictions might have been during the course of a game. We validated that both of these metrics corresponded strongly with those from an expert sports analyst for both positive and negative memories of games (Figure S6A-B). We also validated that full-game surprise from our algorithm corresponded strongly with full-game surprise based on pre-game betting odds (Figure S6C).

**Figure 3:**
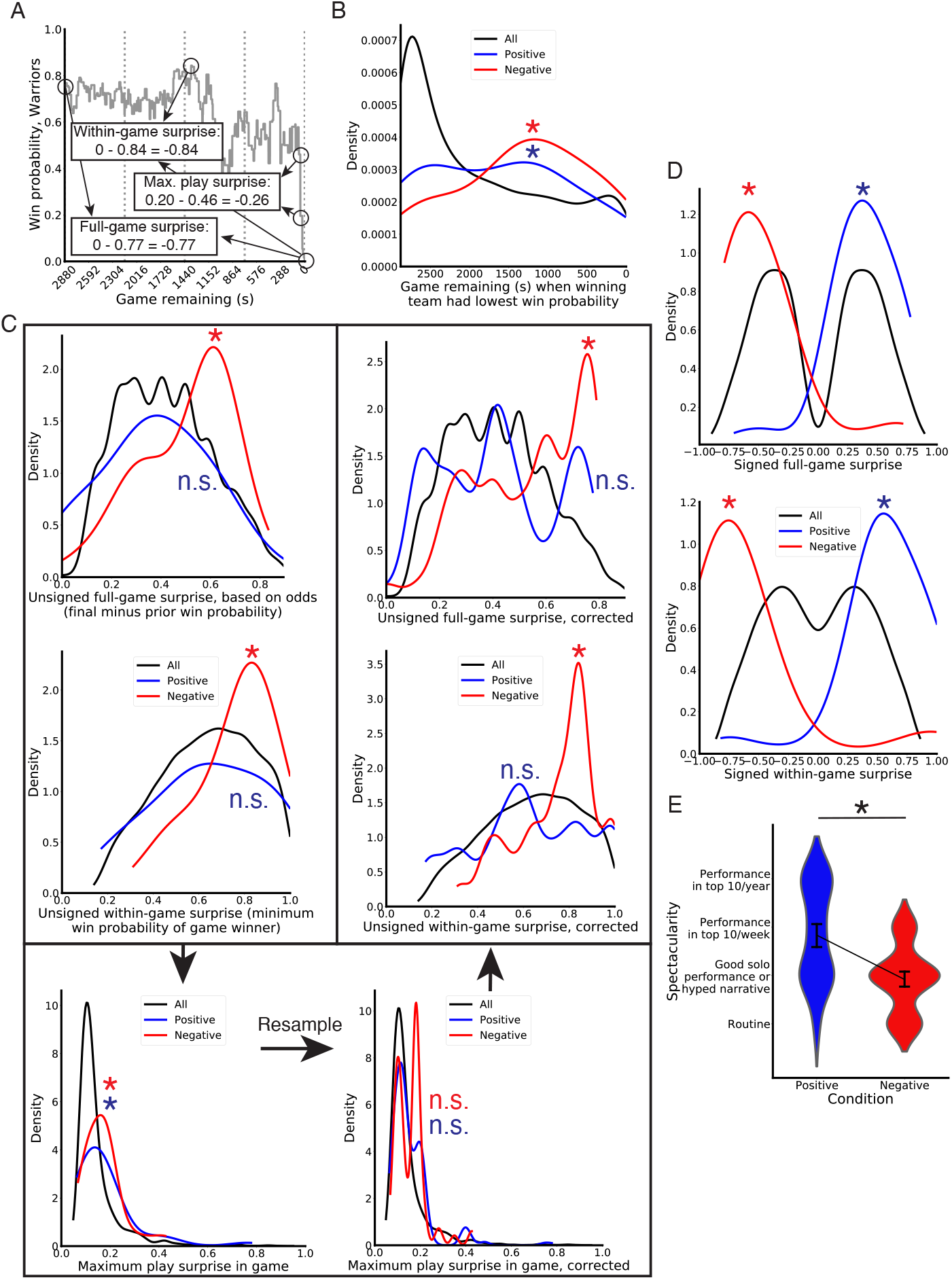
Full- and within-game surprises predicts memory for positive and negative games. (A) We computed full-game surprise as the difference between the pre-game and final win probability and within-game (“comeback”) surprise as the difference between the lowest and final win probability for the game winner. Additionally, we found the maximum play surprise for each game. (B) The game time from when the winning team had their lowest win probability, marking the onset of a comeback, was later in the game for negative, but not positive games. These times were not heavily distributed towards the end of the game, which suggests there is more than a single play that drove the comeback and therefore it occurred over a longer timescale. (C) Unsigned full-game surprise (top left) and within-game (middle left) were significantly greater than the null distribution (all games) for negative, but not positive, game responses. Maximum unsigned play surprise within a game was also, greater for the positive and negative games chosen by subjects than a null distribution of all games (bottom left). We therefore next selected subsets of the positive and negative games that did not differ from the null distribution (bottom right). Using these modified distributions, negative games still had greater unsigned full-game (top right) and within-game surprise (middle right) values than the null distribution. (D) Signed full-game surprise (top) and within-game (bottom) values were significantly higher and lower for positive and negative games, respectively, than the null distribution using all games. (E) The spectacularity of a given game was higher for positive than negative game responses. *: *p* < 0.02

Similar to our characterizations for play memories, we showed that both positive and negative game memories were biased towards the end of a team’s season; similar to the temporal distributions of play memories, negative games were more likely to be recalled closer to the end of a team’s season [median percentile, positive: 98.8th, based on *N* = 77; negative: 100th, based on *N* = 74; Mann-Whitney *U* = 2225, *p* = 0.01] (Figure S6D). We also found the time that the winning team had its lowest win probability. We compared these times to those from a null distribution of all games, which tended to peak early with a skewed distribution towards the end of the games (Figure 3B), again reflecting how many game outcomes were nearly certain from an early time point. For both positive and negative game memories, these times occurred later in the game than in the null distribution [median game remaining, positive: 1364 s, based on *N* = 77; negative: 1364 s, based on *N* = 74; null: 2081, based on *N* = 22539; positive vs. null Mann-Whitney *U* = 9.9*10^5^, *p* < 0.001; negative vs. null Mann-Whitney *U* = 1.1*10^6^, *p* < 0.001], showing that the preferred team had a better chance of winning than is typical at some point into the game. However, these low win probability time points came later in the game for negative than positive games [positive vs. negative Mann-Whitney *U* = 3331, *p* < 0.001]. Notably, these times for negative games were not heavily clustered around the final moments of the game, which suggests that negative game memories were not solely driven by games that were lost in the final seconds. Rather, the events in the game leading to the loss were spread across many accumulated – albeit small – surprises over time.

We again asked whether game memories were influenced by unsigned surprise – in this case, surprise aggregated over multiple events spanning a timescale of minutes to hours. Similar to our play analyses, we compared full-game and within-game unsigned surprise values for positive and negative game memories against a null distribution of these values in all games (Figure 3C, top left). The null distribution was biased towards surprise values *<* 0.5 (median = 0.41), as one should expect if pre-game probabilities hold predictive power. Intriguingly, full-game surprises from negative game memories were significantly greater than those from the null distribution and the positive game memory distribution, whereas positive memories were not significantly different from the null [median full-game surprise, positive: 0.41; negative: 0.60; positive vs. null Mann-Whitney *U* = 8.7*10^5^, *p* = 0.998; negative vs. null Mann-Whitney *U* = 5.5*10^5^, *p* < 0.001; positive vs. negative Mann-Whitney *U* = 1980, *p* = 0.001]. To independently assess full-game surprise using a more well-established source, we also computed it based on pre-game Vegas betting odds, and we found similar results [median full-game surprise, null: 0.42; positive vs. null Mann-Whitney *U* = 6.8*10^5^, *p* = 0.86; negative vs. null Mann-Whitney *U* = 4.3*10^5^, *p* < 0.001]. These results suggest that negative game memories preferentially came from instances in which the subject’s preferred team was expected to win. Next, we compared within-game surprise for all games against positively and negatively remembered games. In the null distribution, these values were above 0.50 (mean = 0.64, median = 0.65) (Figure 3C, middle left). To explain this, consider that scoring in a basketball game is a stochastic process, and simulating a basketball game between two equally strong teams resembles examining whether heads occur more than tails at a coin flip game over 100 flips; the eventual winner between heads and tails is likely to be losing at some point during the coin flip game, leading to > 0.50 within-game surprise on average. Positively remembered games were no different than the null distribution of all games, whereas negatively remembered games had significantly higher within-game surprise than the null distribution and the positive memory distribution [median within-game surprise, positive: 0.62; negative: 0.84; positive vs. null Mann-Whitney *U* = 8.3*10^5^, *p* = 0.556; negative vs. null Mann-Whitney *U* = 5.4*10^5^, *p* < 0.001; positive vs. negative Mann-Whitney *U* = 2055, *p* = 0.003]. These results suggest that, similar to results for full-game surprise, the preferred team was winning by a larger-than-average value at some point during negatively remembered games (i.e., they blew the lead).

We next examined an alternative hypothesis: since games are composed of plays, negative game memories could simply be driven by single plays rather than the more long-term surprise of multiple accumulated events that are by themselves not particularly surprising. The distribution of times from which the lowest win probability occurred was not clustered near the end of the game (when the largest surprises tend to occur) (Figure 3B), which argues against the case for single plays driving game memories. Nevertheless, to further address this concern, we found the most surprising play in the positive and negative games (Figure 3A), and we compared this value to a distribution of maximum play surprise values for all games. As the distribution of maximum surprise values for each game rather than all values, this null distribution is higher than the play surprise distribution (median: 0.118). Importantly, both positive and negative maximum play surprise distributions were higher than the null distribution and did not differ [median maximum play surprise, positive: 0.134; negative: 0.179; positive vs. null Mann-Whitney *U* = 6.9*10^5^, *p* = 0.002; negative vs. null Mann-Whitney *U* = 5.8*10^5^, *p* < 0.001; positive vs. negative Mann-Whitney *U* = 2660, *p* = 0.48] (Figure 3C, bottom left), potentially leaving this as an alternative account for the full- and within-game surprise results. To rule out this account, we first resampled our data with replacement until we found 100 instances in which each maximum play surprise distribution did not differ from the null (Figure 3C, bottom right) (*p >* 0.10). We then tested whether these subsets of games still showed differential levels of full-game and within-game unsigned surprise. Negative game memories within this subset had significantly higher full-(Figure 3C, top right) and within-game unsigned surprise (Figure 3C, middle right) than the null distribution in 99% and 100% of the samples, respectively (see Figure S7 for distributions averaged across samples). Positive game memories did not differ from the null, resulting in significant differences in 10% and 7% of the samples. Therefore, while large momentary surprises may partially contribute to memories for events aggregated across numerous moments like games, they cannot fully explain them, because the effect of game surprise persisted after controlling for play surprise. This indicates that subjects preferentially remembered negative games with surprise aggregating over multiple events.

We next asked whether positive and negative game memories were drawn from games in which subjects’ preferred teams overperformed or underperformed. We again created a null distribution by considering full- and within-game surprise values as if a hypothetical fan cheered once for each team per game. Full-game signed surprise for positive and negative game memories was significantly higher and lower than this null distribution, respectively (Figure 3D, top) [median, positive: 0.41, negative: -0.54, null: 0.0; positive vs. null Mann-Whitney *U* = 9.9*10^5^, *p* < 0.001; negative vs. null Mann-Whitney *U* = 2.6*10^6^, *p* < 0.001; positive vs. null Mann-Whitney *U* = 5245, *p* < 0.001]. We found similar results for within-game surprise (Figure 3D, bottom) [median, positive: 0.55, negative: -0.74, null: 0.0; positive vs. null Mann-Whitney *U* = 5.3*10^5^, *p* < 0.001; negative vs. null Mann-Whitney *U* = 2.9*10^6^, *p* < 0.001; positive vs. null Mann-Whitney *U* = 5188, *p* < 0.001]. As positive game memories did not have higher unsigned full- and within-game surprise values than the null distribution, this analysis supports the idea that these games were good for their preferred team but not especially surprising overall. Conversely, negative game memories had higher unsigned surprise than expected, and these surprises also came with games in which the preferred team underperformed expectations.

Finally, similar to how one might consider a spectacular play to be surprising in an alternative manner to our win probability-derived metric, we also independently rated the spectacularity of games against a hypothetical corpus of other games. We operationalized game spectacularity as (1) a routine game where nothing especially notable occurred, (2) a good solo performance or strong media narrative leading up to the game (e.g., the final game between two historic players), (3) a top-10 performance by a particular player in a given week, or (4) a top-10 performance by a player in a given year. An example positive game memory along these lines reported by 6 subjects was for January 23, 2015, when Golden State Warriors player, Klay Thompson, broke a record by scoring 37 points in a single quarter. We found that spectacularity for games, like for plays, was significantly higher for positive than negative memories [positive: median = 3, mean = 2.8, *N* = 61; negative: median = 2, mean = 1.9, *N* = 74; KS statistic = 0.36, *p* < 0.001] (Figure 3E).

### Long-term surprise influenced memories for positive and negative seasons

To assess whether memories could be influenced by prediction errors across far longer intervals, such as months, we assessed positive and negative subject memories of entire seasons. This analysis involved a different approach, using sports betting odds to model prediction rather than via derivations from play-by-play data. Sports betting odds are released at multiple time points throughout a season, which bettors can use to place money on which team will win the championship. We used these odds to predict final season outcomes for all NBA teams and examined how positive and negative memories for seasons unfolded relative to these predictions (Figure 4A). To assess final season outcomes, we devised a novel system that ordinally ranked teams according to how long they lasted into the season and postseason, and we broke ties using their regular season records (see Methods for more details). Under this system, the championship winner was assigned #1 and the team with the worst record was assigned #30 (Figure 4B).

**Figure 4:**
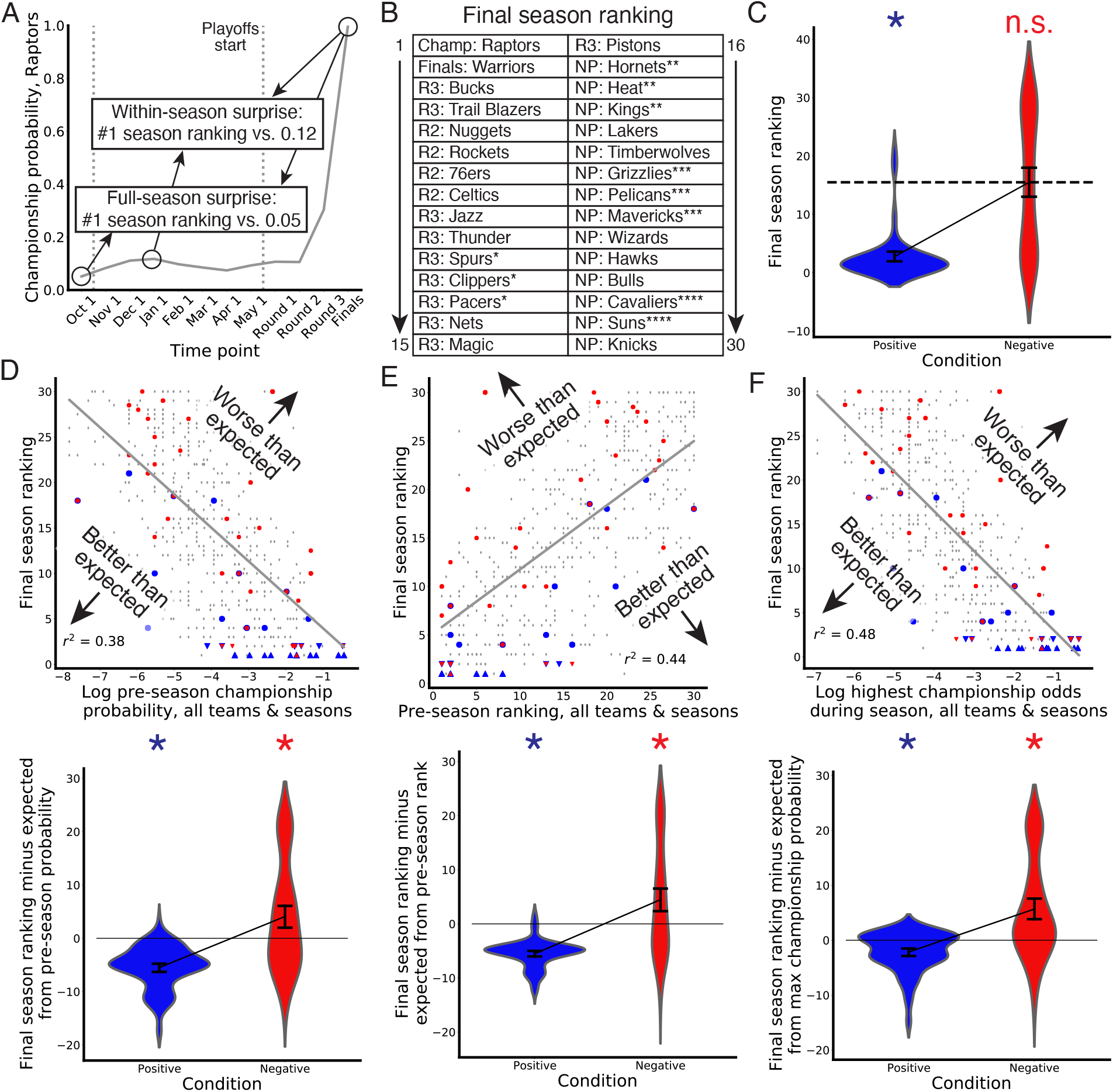
Surprise within and across full seasons predicts positive and negative season memories. (A-B) To calculate surprise across seasons, we first found Vegas championship betting odds updated monthly and before and after each playoff round, shown in (A) for the 2018-19 NBA champion Toronto Raptors. We then computed a novel metric which ranks teams by their success in the playoffs and regular season record. This metric was shown in (B) for all teams at the end of the 2018-19 season. (C) Positively recalled seasons were strongly influenced by this final ranking, as subjects chose seasons in which their preferred team was higher ranked than the average ranking (dotted line), whereas negative seasons did not differ from average. (D-F) To assess how season memories may have differed based on how outcomes deviated from predictions, we used linear regression to predict final season rankings based on the following (top graphs): pre-season championship probability (D), pre-season ranking (E), and highest within-regular season championship probability (E). In each top plot, all seasons from all teams were shown as gray diamonds, positive season memories blue circles, and negative season memories red circles, with championship winners and runner-ups marked instead as upward and downward arrows, respectively. Regression lines were shown in gray. In each bottom plot, we plotted the residual between final outcomes and expectations based on each regression. Positive and negative season memories were better and worse than these expectations, respectively, for each residual metric. *: *p* < 0.01

First, we asked whether final outcomes influenced memories apart from predictions. In this case, we used Wilcoxon signed-rank tests with the average final season ranking as the population mean (15.5) rather than comparing these results against the null distribution of a ranking system, which is a uniform distribution. We found that positive season memories had significantly better rankings for the subject’s preferred team than the average season [median = 1, mean = 2.8, *N* = 96; Wilcoxon signed-rank *Z* = 8.6, *p* = 0.001], whereas negative seasons did not differ from the average [median = 16, mean = 15.5, *N* = 83; Wilcoxon signed-rank *Z* = 0.89, *p* = 0.38] (Figure 4C). To assess the role of prediction in season memories, we first measured the linear relationship for the final season outcomes from two variables constituting full-season surprise and one variable constituting within-season surprise. These variables were the logs of pre-season championship probabilities, pre-season rankings (which ranked teams ordinally according to the pre-season probabilities), and the logs of the highest within-season championship probabilities (excluding the playoffs). All three metrics predicted final season outcome (Figure 4D-F, top) [*r*^*2*^ in predicting final rankings, log pre-season odds: 0.38; pre-season rankings: 0.44; log highest within-season probability: 0.48, all *p* < 0.001]. Next, we used the residuals from these linear relationships to assess how positive and negative season memories measured up in comparison to what was predicted before and during the season. For all three metrics, positive season memories were better than predicted [log pre-season odds: median = -4.8, mean = -5.5, Wilcoxon signed-rank *Z* = 8.4, *p* < 0.001; pre-season ranking: median = -4.8, mean = -5.5, Wilcoxon signed-rank *Z* = 8.5, *p* < 0.001; log highest within-season odds: median = -1.7, mean = -2.1, Wilcoxon signed-rank *Z* = 4.9, *p* < 0.001] and negative memories were worse than predicted [log pre-season odds: median = 1.4, mean = 4.0, Wilcoxon signed-rank *Z* = 2.8, *p* = 0.005; pre-season ranking: median = 2.6, mean = 4.5, Wilcoxon signed-rank *Z* = 2.8, *p* < 0.004; log highest within-season odds: median = 1.8, mean = 5.7, Wilcoxon signed-rank *Z* = 5.1, *p* < 0.001] (Figure 4D-F, bottom).

We next examined whether our results were driven by subjects reporting won or lost championships for their favorite team, which may play a special role for sports fans beyond simply winning games^63^. Therefore, we re-ran these analyses excluding both finals teams in the championship game. In all cases, positive and negative seasons were better and worse than expected, respectively (all *p* < 0.003).

Next, just as numerous plays compose a game, numerous plays and games compose a season. We therefore asked whether these data could be alternatively explained by especially surprising plays or games. For this analysis, we found the largest play surprise, full-game surprise, and within-game surprise values pooled across the entire season for each team in the corpus and for the teams represented by positive and negative season memories. Maximum play unsigned surprise across the season did not differ between the null distribution of all team seasons and those from positive and negative season memories [median, positive: 0.6, negative: 0.59, null: 0.58; positive vs. null Mann-Whitney *U* = 3.5*10^4^, *p* = 0.72; negative vs. null Mann-Whitney *U* = 2.2*10^4^, *p* = 0.33]. Maximum full-game unsigned surprise values were higher for both positive and negative season memories than the null distribution [median, positive: 0.87, negative: 0.87, null: 0.84; positive vs. null Mann-Whitney *U* = 1.6*10^4^, *p* < 0.001; negative vs. null Mann-Whitney *U* = 1.5*10^4^, *p* < 0.001], and maximum within-game unsigned surprise values were lower for both positive and negative season memories than the null distribution [median, positive: 0.98, negative: 0.96, null: 0.98; positive vs. null Mann-Whitney *U* = 3.0*10^4^, *p* < 0.001; negative vs. null Mann-Whitney *U* = 2.7*10^4^, *p* < 0.001]. This may have arisen because participants generally chose seasons from teams who performed at the extremes of team strength (Figure 4C), and therefore the potential for larger surprise values was higher. Nevertheless, after resampling 100 subsets of these distributions such that they no longer differed from the null distribution, we still found that positive and negative season memories were better and worse than predicted, respectively (percentage of samples in which *p* values were significant: positive, adjusting for full-game surprise = 100%; negative, adjusting for full-game surprise = 92%; positive, adjusting for within-game surprise = 100%; negative, adjusting for within-game surprise = 93%) (Figure S8). Therefore, surprise aggregated across a full season could not be explained by a single play or game.

Finally, just as with game and play memories, we independently assessed the spectacularity of reported seasons. We operationalized the season spectacularity of a team as (1) a typical season where nothing especially notable occurred, (2) a season in which a remarkable pre-or mid-season event occurred, such as a top-25 player leaving or arriving to the team, (3) a season in which the team was the most discussed across the NBA during the year, such as the 2010-2011 Miami Heat, or (4) an all-time, historic season in terms of media hype or narrative, such as Michael Jordan’s last season with the Chicago Bulls. Contrary to games and seasons, we found that season spectacularity did not differ for positive and negative memories [positive: median = 2, mean = 1.93, *N* = 96; negative: median = 2, mean = 1.97, *N* = 83; KS statistic = 0.032, *p* = 1.0].

Altogether, subjects reported positive and negative season memories – which involved numerous accumulated games and plays over the course of months – that substantially exceeded and fell below long-term predictions, respectively. Importantly, these findings cannot simply be explained by considering championship teams or surprises on a shorter timescale, such as especially surprising plays or games. These results were captured pointedly by one subject annotating their own memory of a relatively, though not objectively, positive season for their favorite team: “The [Sacramento] Kings have been a bad team ever since I can remember, so my emotions correlating to my favorite team have more to do with my experience than with the performance of the team. I don’t think the Kings have even made playoffs since 2002.”

## Discussion

We found that subjects’ reported memories were biased towards events in which surprise unfolded on the scale of seconds, hours, and months. Given that fans enjoy good outcomes and dislike bad outcomes for their teams relative to expectations, as humans show pleasure or displeasure relative to expectations in a range of domains^90–93^, one might expect that subjects would recall disproportionately high positive and negative *signed* surprises. However, reported memories also had disproportionately high *unsigned* surprises, suggesting that reported memories were not simply due to fan preference. Individually surprising moments (in this case, for individual plays) were well-remembered, which replicates prior results on momentary surprise^1–3,7,11,94^ and extends them into real-world domains^64,65^. The latter aspect is important, because extending ideas from laboratory-based experiments in reinforcement learning (which consist of numerous repetitions of nearly identical trials) into real-world contexts (in which situations may not be precisely repeated) has been a central challenge^95^. More intriguingly, surprising games and seasons – which involved surprise accumulated over multiple sub-events and longer timescales – were also well-remembered. Critically, surprises at these longer scales could not be explained by particularly large surprises at shorter scales (plays in the case of games; plays and games in the case of seasons).

We predicted that subjects would preferentially remember games with high full-game or within-game surprise, which relate to the common terms of “upsets” and “comebacks”, respectively. However, this prediction was only supported for negative game memories. We offer three possible reasons for this. First, when responding for positive games compared to negative games, subjects relied more heavily on spectacularity, an alternative form of surprise. This may have led them to de-emphasize win probability-based forms of surprise as a heuristic in reporting their memories. Though this may mask a positive result, future studies with more leading questions may show the importance of this metric (e.g., “What was your most positive and surprising sports memory for an individual game?”). Second, subjects often chose games in which their team won a championship (35 / 77 positive games), which is a primary goal of sports^63^. Often when teams win the championship, their team is already favored to win the final game owing to their superiority, which would reduce both full-game and within-game surprise due to being expected to outscore the other team throughout the game. From a surprise perspective, this result may be remain puzzling, as it is unclear why such an incremental increase in championship likelihood – even to 100% – should be so positive. We speculate that the jump from a near-certainty to a certainty – and all of the surrounding accolades, bragging rights, and victory parades that follow, perhaps after years or decades of unfulfilled fandom – is positive enough and possibly reflects a strong enough *reward magnitude* that fans value these games in spite of the previously high expectations. Factoring in the subjective reward magnitude of winning a championship into an expected value of magnitude times probability could account for this preference^96^.

A final possibility relates to the phenomenon of “wishful thinking” on the part of sports fans: sports fans of a preferred team often provide biased game reports^97,98^ and overestimate their likelihood of winning^61,99^ (for this effect in other contexts, see^56,60,100–102^). We estimated prediction using our objective win probability model, but wishful thinking could bias this prediction: if the subjective prediction were higher than the objective one for the preferred team, the positive surprise for preferred outcomes would be dampened and negative surprise would be exacerbated. This would lead to greater disappointment for sports fans of the losing team^103–105^ and could be partially responsible for the asymmetries we observed in game memories.

In spite of the overall findings consistent with surprise driving long-term memories, many negative season memories exceeded pre-season predictions but corresponded to seasons in which the preferred team lost the championship (season rank #2 in Figure 4C). These “near-misses”^106,107^ have been reported as aversive^108–110^ and increase with a higher perceived likelihood of the preferred outcome^111^. Yet, it is notable that many championship losses were also regarded as positive season memories. We speculate that this may come down to the perspective (or timescale) one takes on such a season, how one realistically values alternatives^112^, and the specific value one places on championships versus other outcomes^63^. In other words, it may rely on whether it was enjoyable to see a team do so well from the vantage point of the beginning of the season or agonizing to see them fail after being so close to being champions.

To explain the long-term surprise effects in this paper, we argue that sports fans must hold some lingering trace of their predictions from prior time points that become compared against much later outcomes. This presents an interesting puzzle for how these memory biases might arise computationally. Though speculative, consider that an experienced sports fan learns the value of game states in a way that resembles a reinforcement learning agent trained via temporal difference learning (hereafter, TD-RL agent). That is, when encountered with a successive series of states before some final reward outcome, a TD-RL agent learns to predict the value of each state based on its history of reward probabilities and magnitudes^41,113^. Similarly, experienced sports fans can learn to associate game states with win probabilities^7^. Next, consider the following scenarios. In one scenario, the agent’s preferred team has a 98% win probability with 10 minutes remaining in a game, and in another scenario, they have a 10% win probability. After this, both cases converge, in that the win probability gradually decreases or increases to 30% with 1 minute remaining. Then, the preferred team loses by the same score in both scenarios. In either scenario, the experienced sports fan, like the TD-RL agent, will have updated their state probabilities (to 30%) by the 1-minute time point, and therefore the final negative signed surprise is equal in both cases. Additionally, with the same final score, the difference between the scenarios cannot be explained by the overall accumulation of positive and negative events. Therefore, in order for long-term surprise to affect memory, we speculate that the sports fan must not simply update their predictions in a memoryless^114,115^ and rapid manner but instead must keep traces of predictions beyond the immediate moment – such as from the 10-minute mark in these scenarios.

It is intriguing to consider which neural processes produce long-term surprise effects. In sporting events, performance differences accrue over time to decide a winner. This arrangement resembles many lab-based evidence accumulation tasks featuring a series of stimuli (e.g., clicks, visual flashes, visual motion) that unfold over time and create a series of states that later become mapped non-linearly onto some outcome variable (e.g., win or lose, left or right reward trial)^116,117^. Recent findings suggest evidence accumulates and affects behavior across multiple events and at multiple timescales in regions across the cortex^118–123^. Therefore, a future study could, for example, examine whether midbrain or cortical areas respond differentially to well-controlled trials that manipulate the long-term, multi-event surprise, such as by manipulating the overall architecture of the trial (e.g., more right clicks at the beginning, creating a 98% probability of a right reward trial, versus more left clicks, creating a 10% probability) while controlling for the overall amount of evidence for each possibility.

Another relevant neural effect follows from findings that neurons and neuronal areas integrate evidence at different time scales^124–129^. In the above 98% vs. 10% scenario, a long timescale representation could be formed with 10 minutes remaining and, because it is not immediately updated due to its long timescale, surprise is higher in the 98% than the 10% case when the final result unfolds. However, whether neural integration – which has generally been studied on the scale of seconds to minutes – operates on scales of months (as in the season data here) remains unclear.

The dopamine system is also relevant to consider. Dopamine neurons respond with respect to deviations from predictions, such that better outcomes are associated with higher responses and worse outcomes with lower responses^43,130^. Additionally, recent studies have shown that dopaminergic regions also respond to unsigned prediction errors^7,14,15^ and changes in beliefs^131,132^. Dopamine neuron responses are modulated by the reward histories of multiple past trials, indicating that they can integrate information beyond the immediately preceding moment to create predictions^133^. However, this integration is distinct from how a momentary response might differ based on multiple accumulated events in the past. Additionally, it has been shown that dopamine neuron responses reflect updated predictions on a nearly moment-by-moment basis, in line with TD learning^134^. As noted above, an overtrained TD-RL agent would make optimal predictions but would not show differential responses based on different history in leading to the same momentary state. Therefore, it is unclear whether dopamine would respond differently in the 98% vs. 10% cases in the final moments and whether it plays a role in these memory effects.

The ability to form long-term memories for arbitrary, non-repeated events is most intimately linked with the function of the hippocampus^135,136^. Regarding the computational approach to this problem, some recent artificial intelligence models have included episodic memory modules that simulate the hippocampus by saving contextual attributes of the current state in an ongoing fashion for later reference^95,137–140^. Here, a reasonable solution to how long-term surprise could influence memory is by maintaining prior predictions by continuously saving them and comparing them against later outcomes. A possible future test of this idea is whether a hippocampal amnesiac patient would differ from controls in having a smaller bias towards long-term surprising information but relatively intact memory for momentary surprising information.

By asking for subjects to report a only single positive and negative memory at each timescale, we showed that surprise affects memory accessibility but not necessarily its availability^75,141^. Probing all or a larger subset of basketball memories could determine whether surprise affects availability as well. Antony et al.^7^ asked basketball fans to freely recall basketball game clips and found that surprise also drove memory for individual plays when subjects were encouraged to respond with everything they could remember. Therefore, this concern has been partially addressed for momentary surprise in laboratory settings, but future studies could examine whether surprise plays a role by probing a broader swath of memories^64,65^. Another promising future direction follows laboratory findings that salient events (such as surprise) boost surrounding memories in some paradigms^24,142,143^, whereas in others they impair them^144^ or have no effect^3,94^ (see^145^ for a short review). In some cases, subjects spontaneously provided the surrounding context for plays, games, or seasons (see Table S1). However, future studies could more rigorously examine whether information before or after surprising events are affected for real-world memories.

The greatest factor in determining whether an event is surprising is the high-level context^146,147^ or schema^148,149^. This factor guides predictions of what should be expected in a particular situation, and therefore surprise varies across domains^150^. Indeed, the broader literature on surprise includes domains as wide-ranging as music^151^, language^51^, politics^64,65^, and narratives^55,148,152,153^. In this paper, we used the term, “surprise”, to refer to changes in game or championship probability. While our data suggest that memories were also influenced by other variables, such as seeing spectacular feats of athleticism (which we assess in our spectacularity measure and argue provides another form of surprise), who wins a game or championship may be the most relevant variable for sports fans. If the primary task or context differed, such that they were asked to instead count the number of passes or listen carefully to the commentary, we would expect other factors to drive memory. Just as a routine play (e.g., a layup) occurring at a critical moment and corresponding to high surprise would be considered highly surprising in our model, an everyday word would be surprising in a particular context (e.g., “Tornado wind speeds reach over 100 dogs per hour.”).

The outcomes of sports contests can be instrumental to one’s sense of identity^67^ and mood^68^, influencing neural substrates heavily linked with emotion^7,89,154^ and physiological factors like testosterone^155^ and creating lasting memories^69^. In fact, our subjects often noted that both positive and negative outcomes were among the best or worst things that happened to them in a given month or year (and in a few cases, even a lifetime) (Fig S9B). Dovetailing with a prior report showing that surprise influences a host of factors related to memory and emotion^7^, these findings underscore the importance of initial predictions in determining how events are remembered in the real world. Our novel approach of probing memories over multiple timescales and events shows that long-term surprises can influence memory in a manner similar to momentary surprise, widening the door for more behavioral and neural questions investigating their underpinnings and highlighting an overlooked influence in studies of prediction and memory.

## Methods

### Subjects

Undergraduate subjects (*N* = 122, 34 female, 18-28 years old, mean: 19.5) were recruited at California Polytechnic State University, San Luis Obispo (Cal Poly). This school is located along the central coast of California and therefore conveniently pools students who grew up as fans from at least four major NBA fan bases (Los Angeles Lakers, Los Angeles Clippers, Sacramento Kings, and the San Francisco Area-based Golden State Warriors). Subjects took the study for course credit among a number of alternative studies. All subjects considered themselves basketball fans and attested to having seen or played in more than 50 basketball games. All subjects gave consent according with Cal Poly IRB #2020-068.

### Questionnaire

Before the study, subjects were screened as basketball fans and were told they would be responding with autobiographical positive and negative memories of plays, games, and seasons. We specifically asked them to recall everything from memory and not to look up any information. Each prompt began with an openended question about the memory, e.g., “As a FAN, what is your MOST POSITIVE memory of a SINGLE PLAY in a college or professional basketball game?” We asked them to describe in as much specific detail as possible what happened on the court (the play, game, or season itself), any contextual information about when it occurred, such as plays, games, or seasons leading up to the chosen memory, and then information about their experience, such as where they were, how they watched it or tended to watch it, and who they were with or tended to watch it with. To ensure we had as many identifying details as possible for plays and games and to provide for as rich an account as possible given the multidimensional nature of memories^156^, we separately asked numerous clarifying follow-up questions. We asked about the teams involved and when the memories occurred, asking subjects to be as specific as possible without looking anything up. We then asked about fandom, including the team for which they were cheering, how big of a fan they were (1 = almost indifferent, 2 = preferred team in that sport, 3 = favorite team in that sport, 4 = favorite team in any sport), whether they were also specifically cheering against the other team (or teams) and how much they disliked that team or teams (1 = indifferent/almost indifferent, 2 = generally root against the team, 3 = least favorite team in that sport, 4 = least favorite team in any sport). Next, we asked about the emotional impact of the memory, including to rate it with respect to other memories in their life. They rated each memory on an 11-point scale (−5 to +5), as the worst/best thing that happened ever/in a given year/month/week/day or that it was a neutral memory (rating of 0). They also estimated how long the play affected them on a 5-point scale, with 1-4 indicating minutes, hours, days, and months, and 5 indicating that they were still affected. Finally, in light of data suggesting that positive sports memories may be revisited more often than negative ones^157,158^ and the general fact that rehearsing information makes it more accessible, we also asked how many times they estimated they rewatched the play (for play memory), the full game or parts of the game (for game memory), or parts of the season (for season memory).

After responding to each of these questions, we also collected data on subjects’ favorite and least favorite men’s and women’s NCAA basketball teams, favorite and least favorite NBA and WNBA teams, their favorite sport, their favorite team in any sport, an estimation of the number of basketball games they had watched (0-50, 50-200, 200-1000, 1000+), whether they had played basketball [in an organized fashion, only recreationally (e.g., pickup games), only a little (e.g., gym class), or never], and if so, how long they had played.

Subjects were given the option to respond as fans of any major college or professional American sports league (men’s and women’s NCAA, NBA, WNBA). While we originally anticipated a more even breakdown of college and professional responses, they skewed heavily towards the NBA in every major category: Of the 122 responses, we were able to identify 70 as positive NBA plays, 77 positive games, 96 positive seasons, 74 negative plays, 74 negative games, and 83 negative seasons. Therefore, we chose to focus our analyses on the NBA for simplicity and to avoid under-powering other analyses, though surprise drives memory in men’s college basketball^7^ and likely other leagues.

### NBA play-by-play data

Play-by-play data from 17 seasons (2004-05 to 2020-21) were first retrieved from [an NBA API](https://github.com/swar/nba_api). These data included information about each game, including the date of the game, the teams involved, quarter and game time updates, score changes, and descriptions of each play. We pared down the data into game events involving a change in context, which included scores, rebounds, turnovers, and tips from jump balls. In total, these data included >5.6M plays from >22K games. We focused on these 17 seasons because nearly all responses were from this time period and they occurred after the NBA expansion from 29 to 30 teams, simplifying the analyses. All geographical moves by teams (e.g., the Seattle Supersonics moving to become the Oklahoma City Thunder) were re-configured to use the current team location and NBA 3-letter code.

### Win probability model

For the main play and game data analyses, we operationalized predictions by creating a win probability metric based on four factors from the play-by-play data: the score difference between the two teams [oriented as positive (negative) when the home team was winning (losing)], the relative strength of the teams, the amount of game time remaining, and the team in possession of the ball^7^. In practice, the first two metrics were combined into one as the ‘expected score difference’ by the end of the game. Using [publicly available data](https://www.basketball-reference.com/), we found for each team the average number of points they scored minus points scored against them during that season; the difference between these subtractions then indicated the relative difference in team strength for an entire game (see^159^ for a similar method using betting odds). Note that this measurement does not capture fluctuations in team strength within a season, whether due to injuries, new player arrivals and departures, or other factors. However, to circumvent this issue, we separately showed that surprise based on publicly available pre-game betting odds also influenced memory (see below). We then divided this by the total number of seconds per game (to estimate how much the stronger team should outscore the weaker team, in units of points per second), multiplied it by the number of seconds left in the game, and subtracted this number from the current score difference. For instance, if for one season the Milwaukee Bucks scored an average of 106 points and had an average of 101 points scored against them (+5 net) and the Boston Celtics scored an average of 98 points and had an average of 97 points scored against them (+1 net), the Bucks would be expected to outscore the Celtics by 4 points over an entire game. Therefore, the expected score difference at the beginning of the game would be 4 points in the Bucks’ favor. If the Celtics led by 1 at halftime, the Bucks would nonetheless be expected to outscore them by 2 points over the course of the second half (4 points per game advantage * 50% of the game remaining), so the expected score difference would be 1 point in the Bucks’ favor.

To create the model, we divided possible game states into a grid in three dimensions: the expected home minus away team score difference from -30 to 30; the number of seconds remaining in the game in increments of 6 (e.g., 2880 – 2774 s, 2774 – 2768 s) until the final 6 seconds, after which we used increments of 2 (e.g., 6 – 4 s, 4 – 2 s, 2 – 0 s); and whether the home team possessed the ball (1/0). Game states from overtime periods were treated identically in the model to those from 4^th^ quarter periods with the same number of seconds remaining^159^. We smoothed the data in the temporal dimension but not the score dimension. For most time points, we combined three time increments (e.g., 2774 - 2758 s), though to avoid edge effects, we combined the first two increments for the first time point (e.g., 2880 - 2768 s). We used finer increments near the end of the game because of the rapid swings in context that occur in NBA games with very little time remaining. Accordingly, we did not temporally smooth the data for the final time bin (2 - 0 s). To compute win probabilities, we found all possible games matching each state and “peeked ahead” to see how often the home team won the game. Probabilities for contexts with a greater difference in score than 30 points were set to 0.998 or 0.002. This model essentially resembles a “look-up table” rather than an algorithm such as logistic regression. We chose this approach due to non-linearities in the state space^159^. For example, win probabilities for the home team in possession of the ball with a -4 to +4 point lead with 5 seconds remaining are the following: 0.018, 0.097, 0.197, 0.36, 0.652, 0.876, 0.938, 0.983, 0.998. Additionally, the win probability benefit of having possession of the ball is not uniform across all score differences and game times but rather peaks when the score is close towards the end of the game (Figure S3)^159^. To validate this approach, training linear and logistic regression models using the same factors yields higher win prediction error rates as defined by mean differences between all predictions and outcomes (mean error for linear regression: 0.355; logistic regression: 0.316; lookup method: 0.304).

Note that our emphasis is not on the process by which subjects learn the value (win probability) of the game states, akin to the goal of a reinforcement learning agent. Rather, we start from the assumption that subjects have predictive mental models of win probabilities that approximate the true probabilities, as previously validated^7^. A rare event – such as a team coming back to win after having only a 1% win probability – does not mean the prediction was wrong if the same game state occurred 99 other times in which the team lost.

### Aligning subject responses with NBA play-by-play data

To find win probabilities and (subsequently) surprise values using our algorithm for play memories, we logged subject responses by the game date, teams involved, quarter, and seconds remaining. Then, we searched through the play-by-play database to find the specific win probability values before and after the chosen play by finding the specific number of seconds remaining when the play ended. In many cases, multiple plays occurred within a single second (such as with free throws, wherein multiple plays elapse with no time expiring, or with shots taken within a single second, such as shots taken with only a fraction of a second on the game clock). In these cases, we manually parsed the data to find the correct moment.

To find win probabilities for game memories, we logged subject responses by the game date and teams involved. To find championship probabilities, we logged subject responses by the season and team involved.

### Surprise metrics

Numerous ways of quantifying surprise have been devised [for a systematic review, see^150^]. All surprise constructs used here were probabilistic point estimates of how outcomes differed from prior predictions. To find play surprise, we computed the difference in win probabilities from before to after the chosen plays. These values were oriented as positive for the home team and were flipped if the subject preferred the away team to correctly align signed surprise. For unsigned play surprise, we instead used the absolute values. To externally validate our surprise metric for plays, we looked up the signed difference in win probability values (signed surprise) given the game context using [an expert analyst's site](http://www.inpredictable.com) (Fig. 2B). This expert used [many of the same factors](http://www.inpredictable.com/2015/02/updated-nba-win-probability-calculator.html) to determine win probability, including game time remaining, score, possession, and team strength (though they used pre-game Vegas odds). They also took special considerations towards the end of the game, during which they abandoned their logistic regression model and opted for a decision tree, resembling our lookup table approach. We compared our surprise metrics against the expert using Pearson correlations.

To find full-game surprise based on our algorithm, we computed the difference in pre-game (prior) win probabilities (with 2880 seconds remaining) and the final probability (1/0). To find within-game surprise, we first found the lowest in-game win probability of the eventual game winner and computed the difference between this and the eventual outcome. For instance, if the home team won and their pre-game win probability was 0.54 and their lowest win probability was 0.22 in the 3rd quarter, full-game surprise would be 1 – 0.54 = 0.46 and within-game surprise would be 1 – 0.22 = 0.78. Similar to play surprise, both of these values were oriented as positive for the home team and flipped if the subject was cheering for the away team. We also computed unsigned values of these metrics using the absolute values. To externally validate our win probability on full- and within-game surprise, we similarly found the pre-game probability and minimum win probability for the winning team on the same expert analyst’s site (Figure S6). As another measure of external validity, we also obtained [pre-game betting odds for each game](https://www.sportsbookreviewsonline.com/scoresoddsarchives/nba/nbaoddsarchives.htm). Odds are posted according to a formula, such that if they are above 0, the championship likelihood is

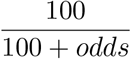

and if they are below zero, the odds are

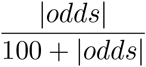

From these probabilities, we computed an alternative measure of full-game surprise. Computing full-game surprise in this way has the advantage of win probabilities being updated more frequently than a full season, which allows for within-season fluctuations in team strength, but it has the disadvantage of coming from a closed algorithm. Nevertheless, full-game surprise values from our algorithm and from the betting odds for the chosen games were highly correlated (positive: *r* = 0.79, negative: *r* = 0.89) (Figure S6), and both ways of measuring full-game surprise produced similar qualitative results.

To find season surprise, we created a final season ranking metric as an outcome variable that used how far teams survived into the season and playoffs to determine ranking, and we split ties using their regular season record. NBA champions were given a #1 ranking, runner-ups were given a #2 ranking, conference championship finalists were given #3-4 rankings (decided by inverse regular season record), conference semifinalists were given #5-8 rankings, etc. If two teams survived the same amount into the season and had the same regular season record, their rankings were averaged (e.g., if #3-4 had the same record, each was given a ranking of 3.5). This metric was more fine-grained than one that is analogous to games, wherein winners are assigned a probability of 1 and losers 0, which would involve giving the champion a 1 and all other teams 0. An alternative metric would be regular season record; however, (1) many fans would consider this unsatisfactory because regular season is not what determines champions, and (2) to prevent injuries in the playoffs, many recent basketball teams have questioned the utility of maximizing their regular season record in favor of simply playing well enough to position themselves for a playoff run while managing workloads^160,161^. Finally, this metric also resembles [how major sports leagues](https://operations.nfl.com/journey-to-the-nfl/the-nfl-draft/the-rules-of-the-draft/) determine the order in which teams can draft new players.

We used three different measures to model subject prediction at the beginning of and throughout the season. Each measure relied on the Vegas championship odds for each team and season as they were updated throughout the year. Before 2009, these odds were only updated before the season and before each round of the playoffs. After 2009, the odds were updated monthly before and during the season and sometimes during the all-star break. We computed the odds in an identical fashion to the pre-game betting odds above. For our first predictor, we used the final pre-season odds for pre-season prediction. Our second predictor resembled the first, except that it ranked teams ordinally by their pre-season odds (and thereby used the same ordinal metric for both predictor and outcome variables). For our third predictor, we used all of the available regular season odds to find the best odds throughout the season for a given team. We used linear regression to find the best fit between these predictors and the final season ranking metric. We used the logarithm of the 1^st^ and 3^rd^ predictors because doing so produced a more linear fit. We then used the residual from these regression lines to determine whether teams did better or worse than expected given this predictive relationship.

### Spectacularity metrics

We devised a spectacularity metric scale from 1-4 meant to assess spectacular or otherwise notable and rare plays, games, and seasons. See main text for descriptions of each rubric.

### Statistical analyses

Our main comparisons in this paper were of three main varieties. First, we compared attributes of positive and negative reported memories against null distributions from the NBA play-by-play corpus and betting odds. In cases in which a shorter time scale form of surprise also differed from the null distribution for a given memory (e.g., maximum play surprise for game memories), we re-sampled the data with replacement 100 times to find instances in which the shorter time scale effect was no longer significant and re-investigated the effect of interest. Second, we directly compared attributes of positive and negative reported memories themselves. For each test, we first conducted skewness and kurtosis tests of normality. In every case, these tests violated assumptions of normality, so we used two-sided Mann-Whitney *U* tests for independent-sample comparisons and Wilcoxon signed-rank tests for one-sample comparisons. Third, we compared distributions of positive and negative responses on various non-surprise attributes, such as spectacularity. These comparisons used two-sample Kolmogorov-Smirnov (KS) tests.

The analyses in this study were not pre-registered.

### Data and code availability

All relevant data and code necessary to reproduce these results, including how to obtain NBA API data, are available [here](https://github.com/JamesWardAntony/bball_am).

## Author Contributions

J.W.A. conceived, programmed, and analyzed data from the experiment, and drafted the manuscript. J.V.D. and J.R.M. contributed to study design and scored a large portion of the recall data. A.J.B. contributed to study design. K.A.B. contributed to study design and coordinated all data collection.

## Acknowledgements

The following individuals assisted with data collection and scoring: Jesús V. Figueroa, Trevor Guerra, Jake Henige, Nami Saito, and Michela Smith. The authors also thank Chung Hee Bennion for her assistance and support throughout the entire project.

## Supplementary tables and figures

**Table S1:**
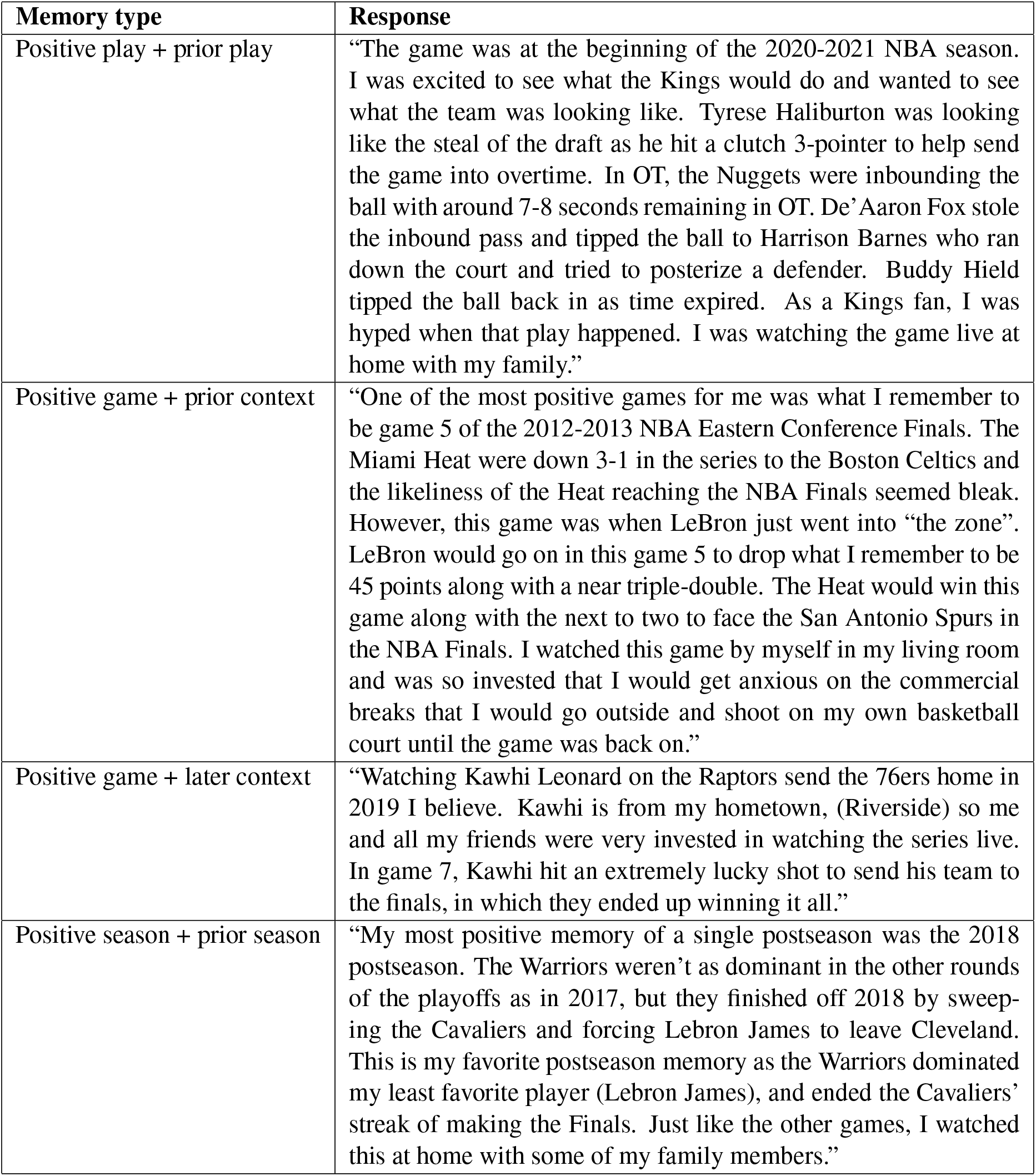
Sample memories in which subjects included the preceding or following play, game, or season context in their description.

**Figure S1:**
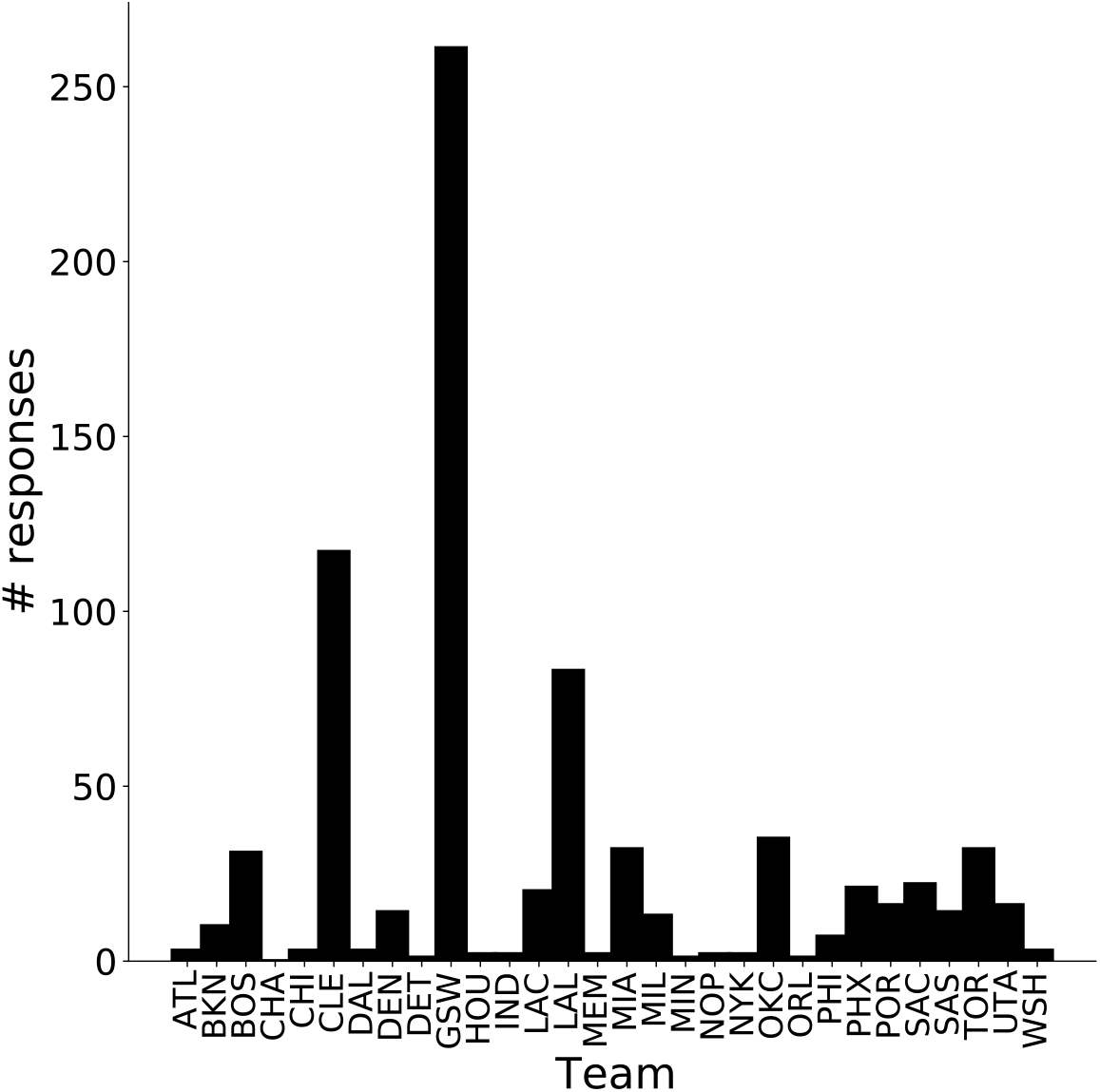
Distribution of responses for each team combined for plays, games, and seasons.

**Figure S2:**
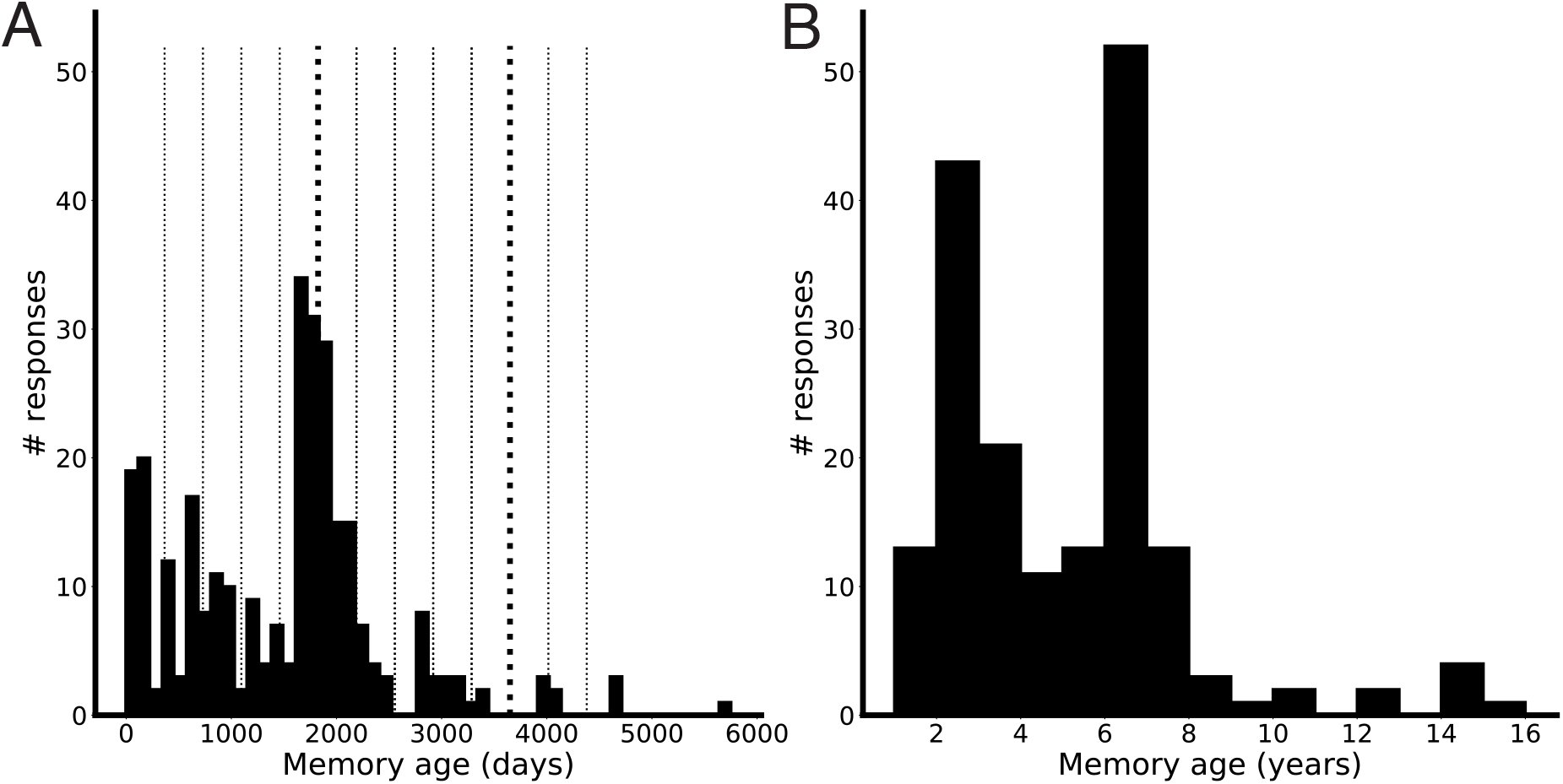
Memory ages. Data were collected over the course of 9 months, so as an alternative to Fig 1C-D, we plotted the time in days between the game date and study date for plays and games (A) or the time in years between the season start and study date (B).

**Figure S3:**
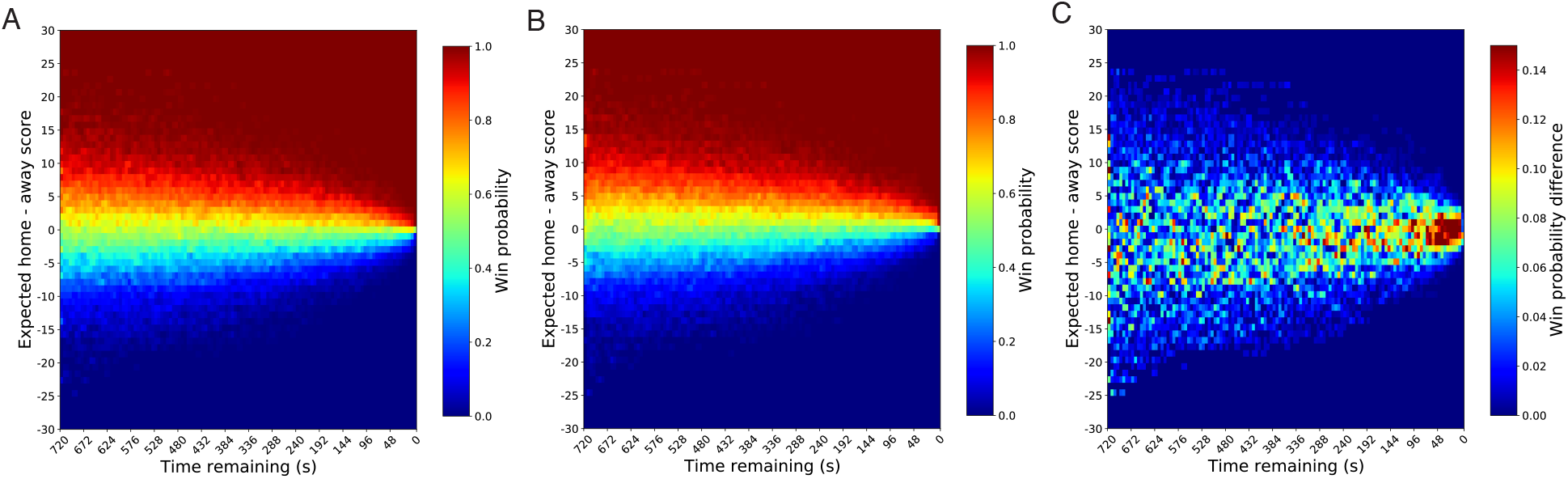
Full win probability model. (A-B) We plotted win probability by expected home minus away score difference and game time remaining when the home (A) and away (B) teams had possession of the ball. To illustrate the importance of team possession, we also plotted the contrast of home possession minus away possession (C). All graphs show data only from the 4^th^ quarter of games.

**Figure S4:**
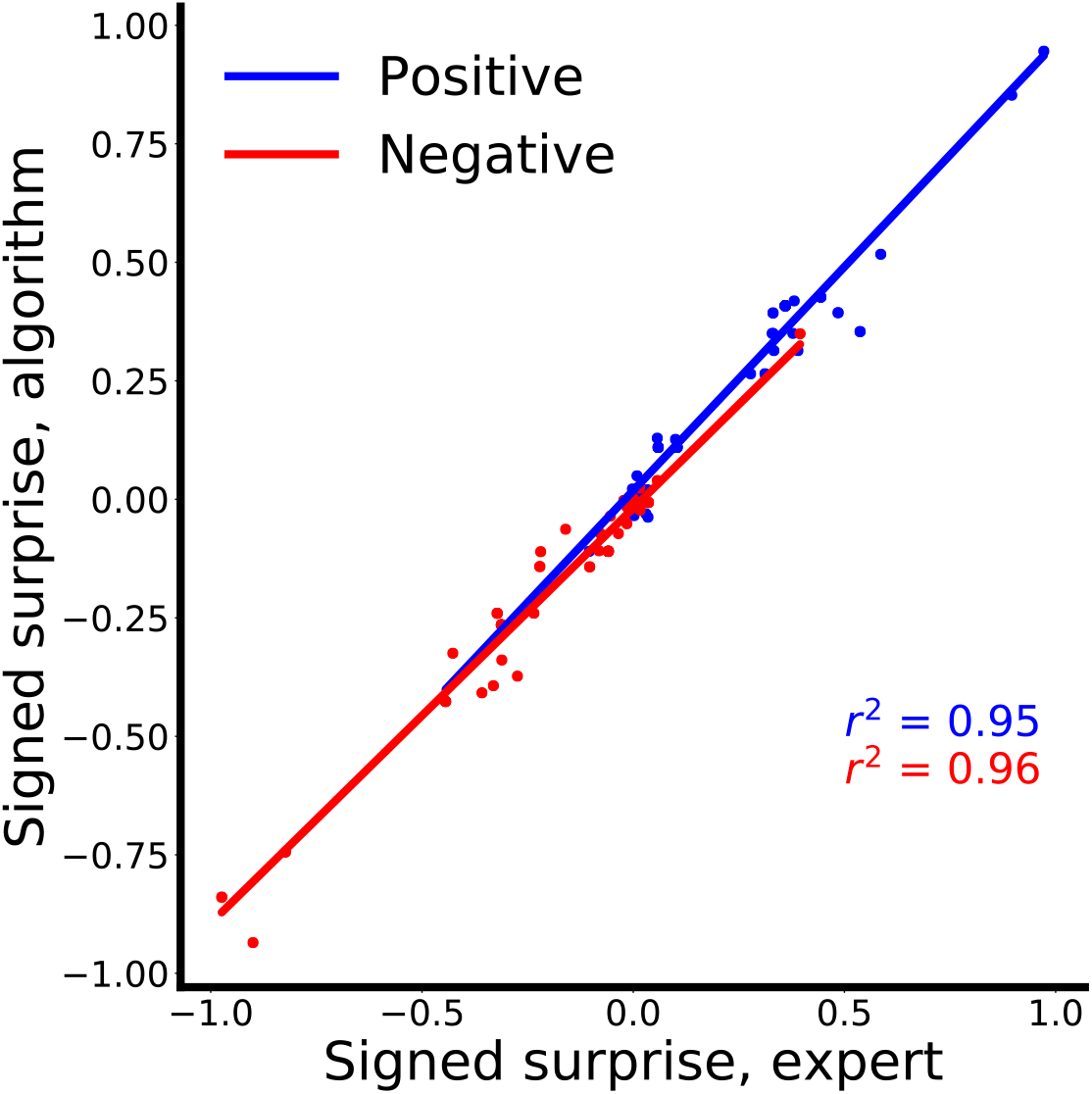
Play surprise values from our algorithm correlated strongly with those from an expert.

**Figure S5:**
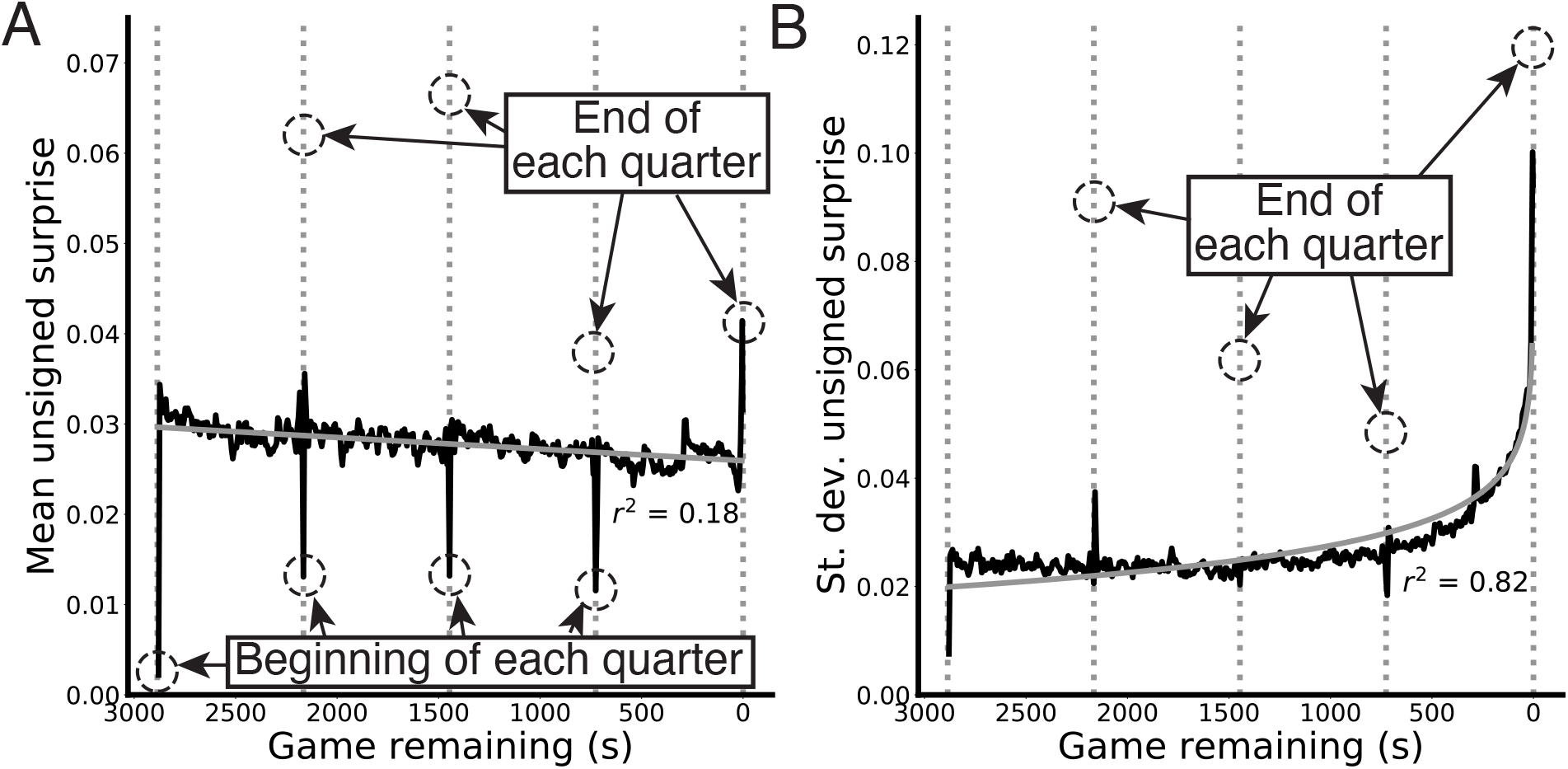
Unsigned surprise over time across a game. (A-B) We plotted the mean (A) and standard deviation of unsigned surprise (B) against game time using our metric. Mean surprise had momentary troughs and peaks at the beginning and end of each game, likely due to the momentary decrease and increase in shots taken within these short time frames. Interestingly, mean surprise tended to decrease throughout the game, owing to the probability that the outcome of many basketball games would be nearly decided at that point. However, standard deviation increased exponentially until the end of games, due to the combination of games with a near-certain outcome, which have almost no surprise, and close games, which involve increasingly rapid swings in win probability space as one nears the finish.

**Figure S6:**
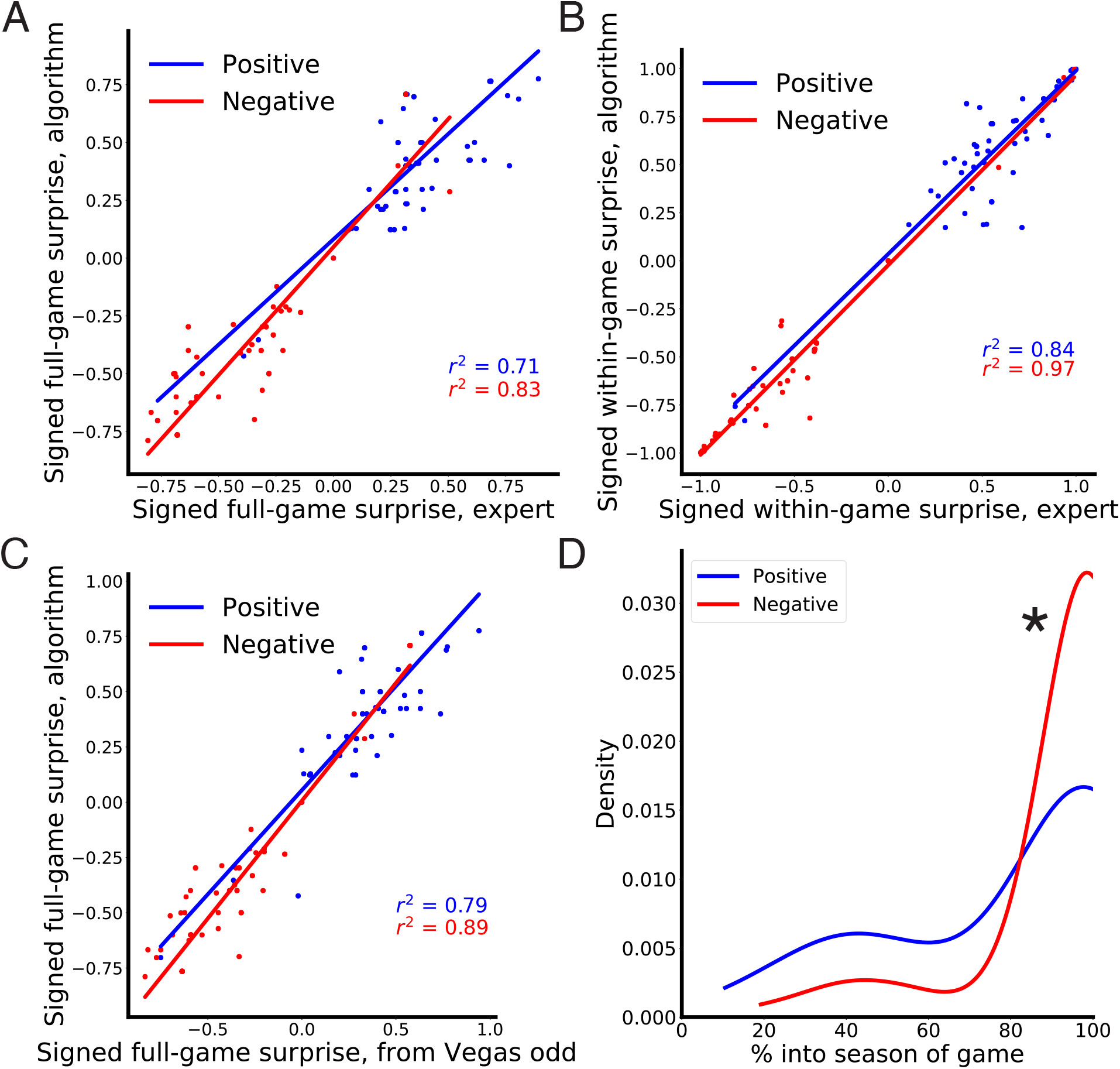
Supplemental information related to game memories. Signed full-game (A) and within-game surprise (B) values correlated strongly with those from an expert site. Similarly, full-game surprise values from the algorithm correlated strongly with those based on pre-game Las Vegas betting odds data (C). Both positive and negative game responses were strongly biased towards the end of the corresponding team’s season, and negative responses were significantly more biased (D). *: *p* = 0.01

**Figure S7:**
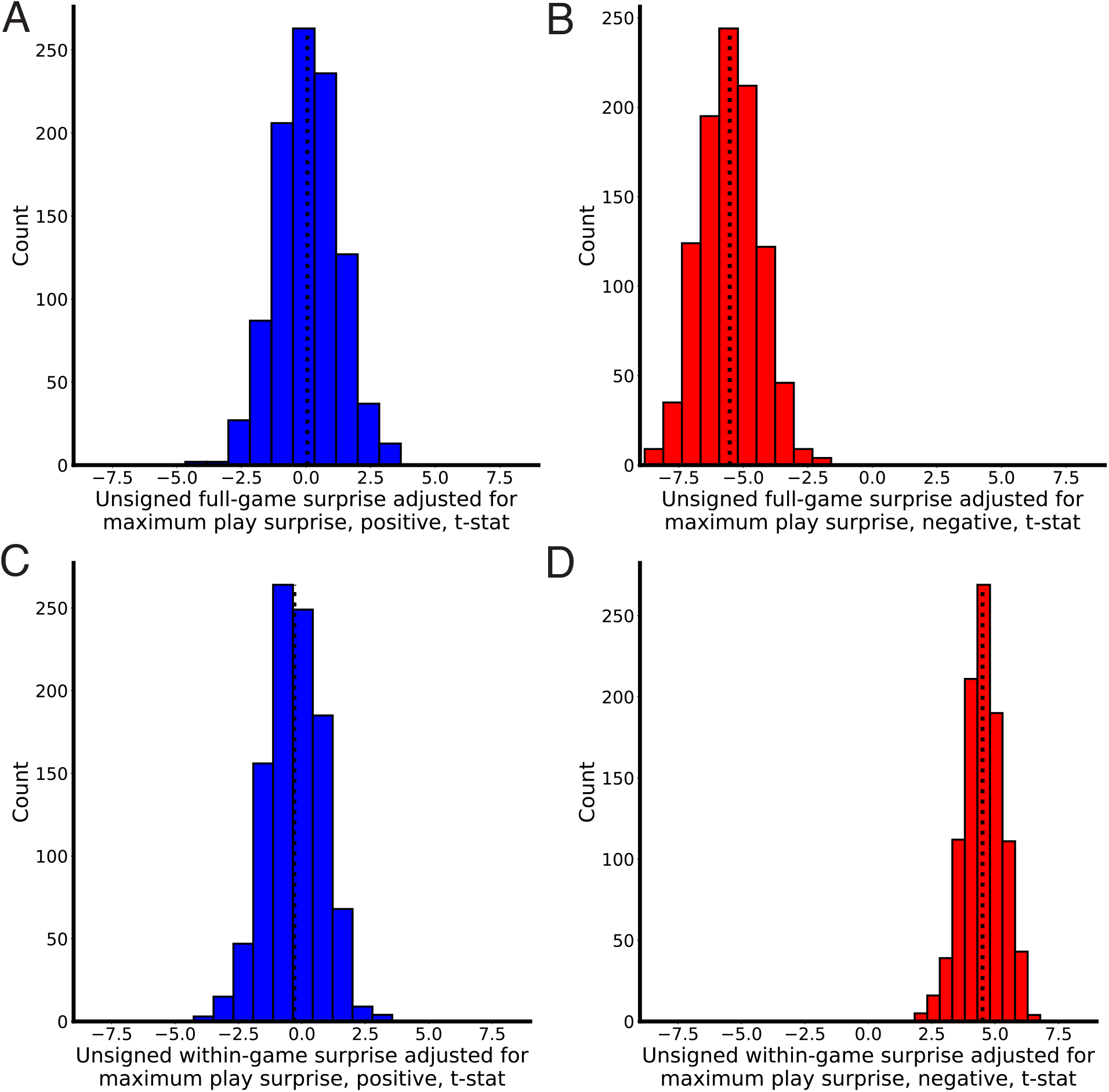
Unsigned full- and within-game surprise after adjusting for differences in play surprise. (A-D) We plotted the *t*-statistics of subsets of the game data that did not differ in maximum play surprise. (A-B) *t*-statistics for positive (A) and negative (B) full-game surprise. (C-D) *t*-statistics for positive (C) and negative (D) within-game surprise. Both negative distributions remained significant, whereas both positive ones did not.

**Figure S8:**
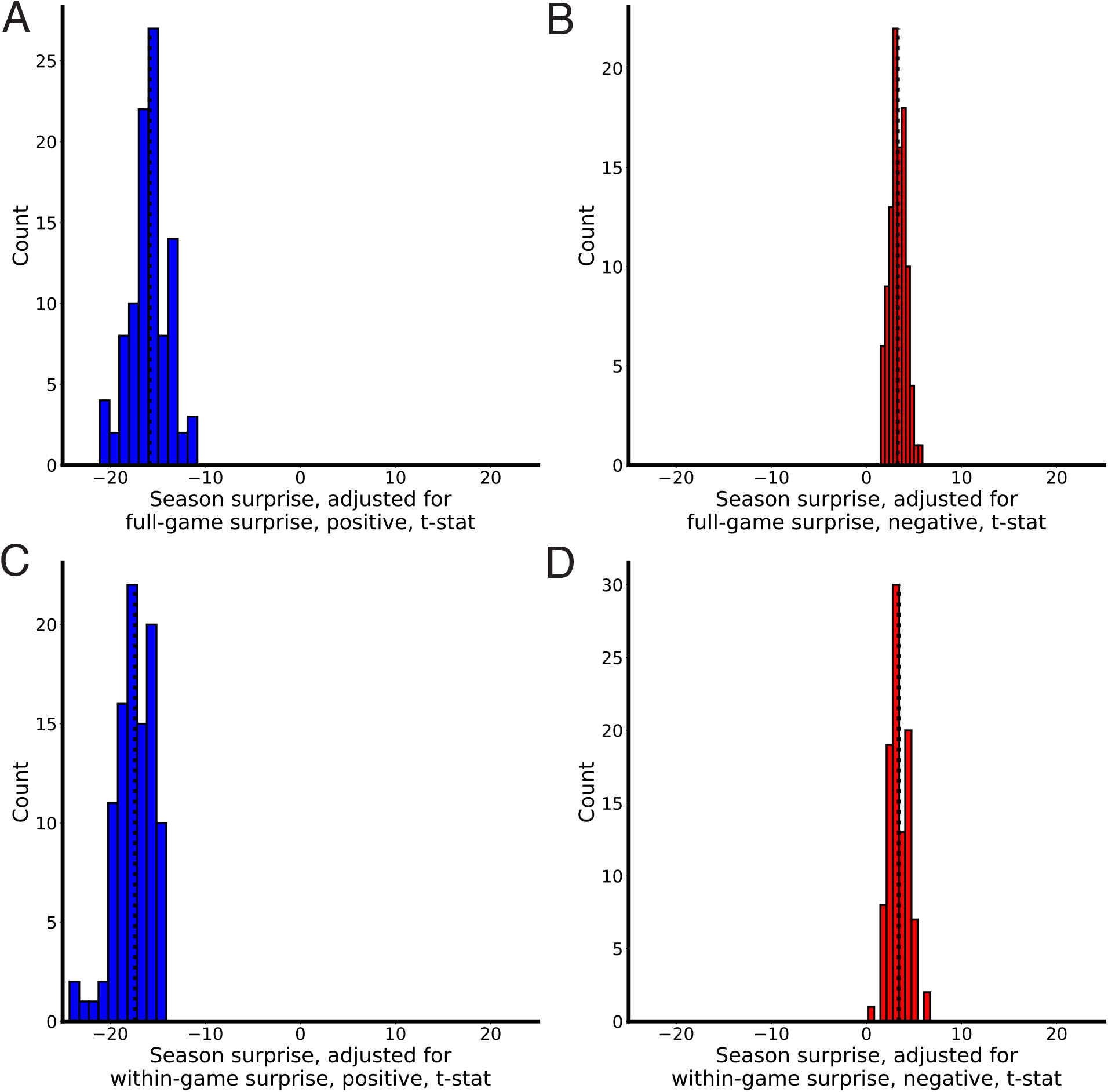
Season surprise after adjusting for differences in maximum full- and within-game surprise. (A-D) We plotted the *t*-statistics of subsets of the season data that did not differ from the null distribution in maximum game surprise. (A-B) *t*-statistics for positive (A) and negative (B) seasons, adjusting for maximum full-game surprise. (C-D) *t*-statistics for positive (C) and negative (D) seasons, adjusting for within-game maximum surprise. All distributions remained significant after adjusting for game surprise deviations from the null.

**Figure S9:**
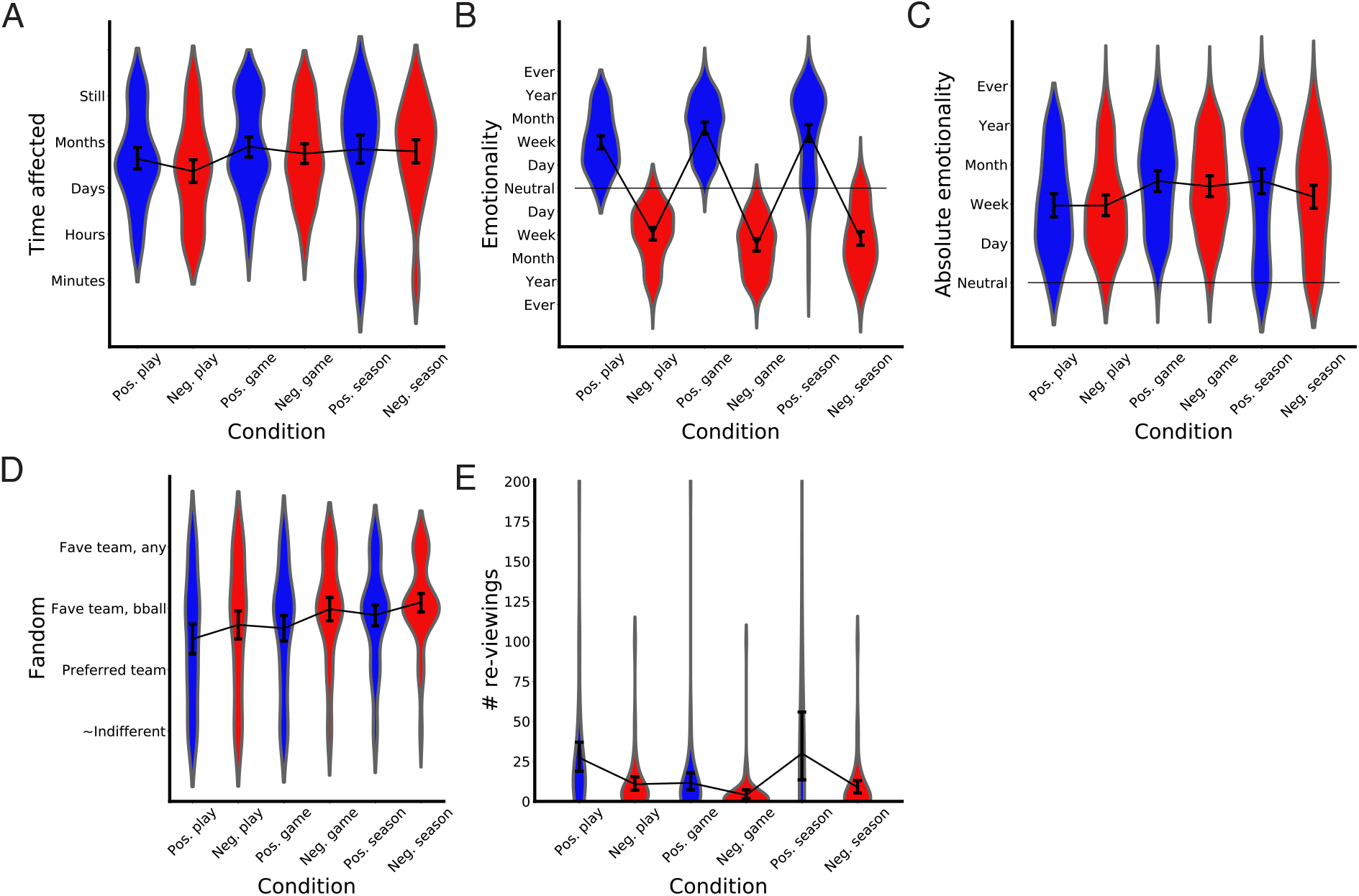
Characteristics of subject memories. We plotted metrics for each type of memory on (A) how long fans were affected by the outcome (1=minutes, 2=hours, 3=days, 4=months, 5=still affected), (B) emotionality with respect to everyday life (−5 to 5; worst to best thing that happened on a given day, week, month, year, ever), (C) unsigned emotionality, (D) their level of fandom (1=almost indifferent, 2=preferred team in that sport, 3=favorite team in that sport, 4=favorite team in any sport), and (E) approximate number of times they re-watched the play, parts of the game, or parts of the season.

